# *Arg1*^+^ microglia are critical for shaping cognition in female mice

**DOI:** 10.1101/2021.08.15.456225

**Authors:** Vassilis Stratoulias, Rocío Ruiz, Shigeaki Kanatani, Ahmed M. Osman, Jose A. Armengol, Antonio Rodríguez-Moreno, Adriana-Natalia Murgoci, Irene García-Domínguez, Lily Keane, Guillermo Vázquez-Cabrera, Isabel Alonso-Bellido, Nathalie Vernoux, Dario Tejera, Kathleen Grabert, Mathilde Cheray, Patricia González-Rodríguez, Eva M. Pérez-Villegas, Irene Martinez-Gallego, David Brodin, Javier Avila-Cariño, Mikko Airavaara, Per Uhlén, Michael T. Heneka, Marie-Ève Tremblay, Klas Blomgren, Jose L. Venero, Bertrand Joseph

## Abstract

Diversity within microglia, the resident brain immune cells, is reported. Whether microglial subsets constitute different subtypes with intrinsic properties and unique functions has not been fully elucidated. Here, we describe a microglial subtype characterized by the expression of the enzyme Arginase-1, *i.e. Arg1* ^*+*^microglia, which is found predominantly in the cholinergic neuron-rich forebrain region during early postnatal development. *Arg1*^*+*^ microglia are frequently observed in close apposition to neurons and exhibit a distinctive molecular signature reflecting a reactive profile. Arg1 deficiency in microglia results in impaired dendritic spine maturation in the hippocampus where cholinergic neurons project, and cognitive behavioural deficiencies in female mice. Our results expand on microglia diversity and provide insights into distinctive spatiotemporal functions exerted by microglial subtypes.

## Main

Microglia, the resident immune cells of the central nervous system, fulfil multiple and contrasting functions across development and adulthood, in steady-state conditions, but also in the context of proliferative, traumatic and degenerative diseases. Microglia are commonly regarded as a population of versatile cells that can acquire distinct phenotypes upon exposure to extrinsic cues in their environment. However, recent high throughput genome wide sequencing data, revealed that microglia with different transcriptomic profiles co-exist throughout the lifespan of mouse during both homeostasis and disease-related challenges^1-5^. Whether these subsets constitute different microglial subtypes with intrinsic differences and functional specialization(s) has not been systematically explored^6^. Worth a notice, these studies show that microglia heterogeneity is strikingly high during postnatal development, when brain is growing in size and establishing its neuronal networks^7, 8^. Postnatal life encompasses critical phases of the mammalian brain development. Indeed, whereas the foundation of the brain development begins before birth, the wiring of some of the neuronal networks, in particular those involved in higher cognitive and sensory functions, as well as sex-related behaviors, take place postnatally. During childhood and adolescence, the brain forms and refines complex neuronal networks through synaptogenesis, pruning and myelination^8, 9^. Interestingly, established microglial biological functions offer a striking match to the above described postnatal brain developmental events^10^. As a matter of timing, microglia populate the brain very early during embryogenesis, from a yolk sac origin, and are present when neuronal circuits start to assemble^11^. Furthermore, beyond their immune functions, microglia are reported to modulate the formation of axonal tracks, synaptic reorganization, turnover and activity and to contribute to the maturation of neural circuits^12-17^. In addition, microglia, in particular a subpopulation of *CD11c*^*high*^-microgia expressing large amount of insulin-like growth factor 1 (IGF-1), are regulators of oligodendrocyte differentiation and myelin formation^18^. A further emerging dimension of the multifaceted microglia is that they exhibit sex differences at least as early as early postnatal development in morphology, maturation and functional output^19-22^. Considering the plethora of functions described for microglia in the developing brain and the reported postnatal microglial transcriptional diversity, one could envisage that distinct subsets of microglia are responsible for exerting these various biological functions. Here we report a novel microglial subtype, *Arg1*^+^ microglia. *Arg1*^+^ microglia are morphologically indistinguishable from neighbouring “canonical” microglia, but defined by a distinct transcriptome profile, as we**l** as a unique spatial and temporal distribution and function in the developing brain.

## Results

### *Arg1*^*+*^ microglia coexist along with “canonical” *Arg1*^*-*^ microglia predominantly in the basal forebrain and ventral striatum of unchallenged mouse brain

In the wild-type (WT) and unchallenged mouse brain of both sexes, a subset of ionized calcium binding adaptor molecule 1 (IBA1, also known as AIF1)-labelled microglial cells was identified based on the expression of the enzyme arginase-1 (ARG1) during early postnatal development. Indeed, immunofluorescence analysis of WT mouse brains at postnatal day 10 (P10) and P28, revealed that ARG1-expressing microglia, thereafter referred as *Arg1*^*+*^ microglia, co-exist in close vicinity with “canonical” microglia that do not express ARG1 (Fig. 1a-b). We also identified ARG1^+^/IBA1^-^ cells in the cerebellum and around the lateral ventricles, which morphologically do not resemble microglia and were therefore excluded from this study (Supplementary Fig. 1). We further confirmed the existence of *Arg1*^*+*^ microglia in the YARG reporter mice^23^, which express yellow fluorescent fusion protein inserted downstream of the endogenous stop codon of *Arg1* gene (Supplementary Fig. 2a).

**Fig. 1.**
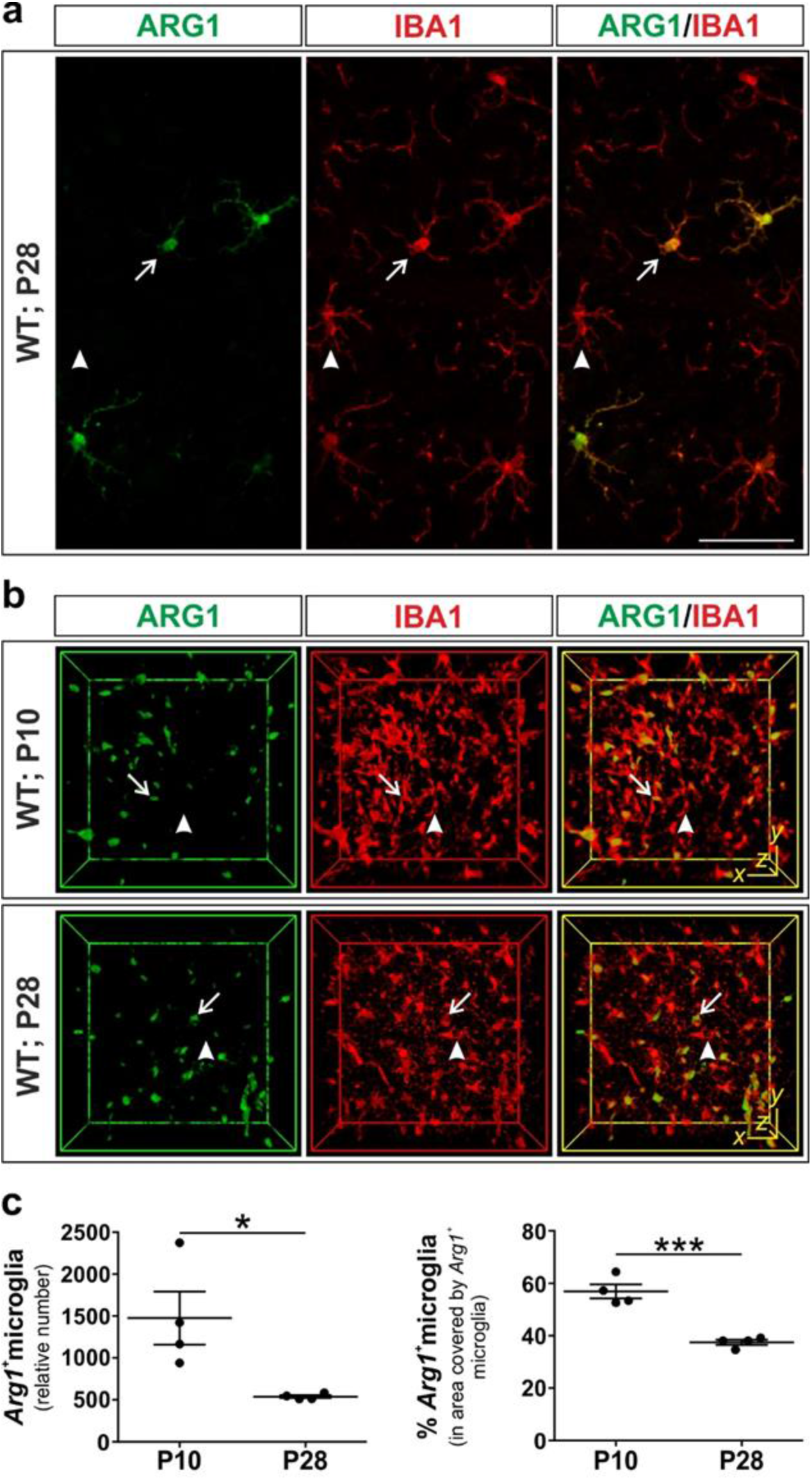
*Arg1*^*+*^ microglia co-exist in the same vicinity with *Arg1*^*-*^ microglia. **a-b**, *Arg1*^*+*^ microglia (arrows) and “canonical” *Arg1*^*-*^ microglia (arrowheads) in WT female (**a**, confocal) and WT male (**b**, iDISCO) mouse brains. **c**, *Arg1*^*+*^ microglia population declines with age (n=4 female animals), both in relative number and in percentage (*Arg1*^*+*^ microglia over total IBA1+ve ce**l** s). Each dot corresponds to one animal. Scale bars, x=50 μm; z=8.25 μm (**A**), x=y=50 μm; z=150 μm (**b**). Data in **c** represented as mean ± s.e.m. Significant differences were determined by an unpaired two-tailed *t* test (\**P*< 0.05, \*\*\**P*≤ 0.001) (**c**).

To gain further insights into the topographical localisation of *Arg1*^*+*^ microglia, iDISCO+ 3D-deep imaging of ARG1 and IBA1 expression was performed on P10 and P28 mouse brains^24^. *Arg1*^*+*^ microglia were found to cluster in several brain regions both at P10 and P28. The *Arg1*^*+*^ microglia located in the basal forebrain (BF) and ventral striatum (vStr) constituted the largest *Arg1*^+^ microglia cluster (Fig. 2a (red cluster), Supplementary Table 1 and Supplementary Movies 1-4). Further registration of the P28 *Arg1*^*+*^ microglia population against the Allen Developing Mouse Brain Atlas (http://mouse.brain-map.org), revealed that the highest concentration of *Arg1*^*+*^ microglia is located in the ventral pallidum (VPal), followed by adjacent areas (Fig. 2b and Supplementary Table 2). The BF is an area rich in cholinergic neurons which project to the hippocampus, a structure engaged in cognition^25, 26^. In fact, loss of BF cholinergic projections and reduction of the BF volume are associated with cognitive decline^25^.

**Fig. 2.**
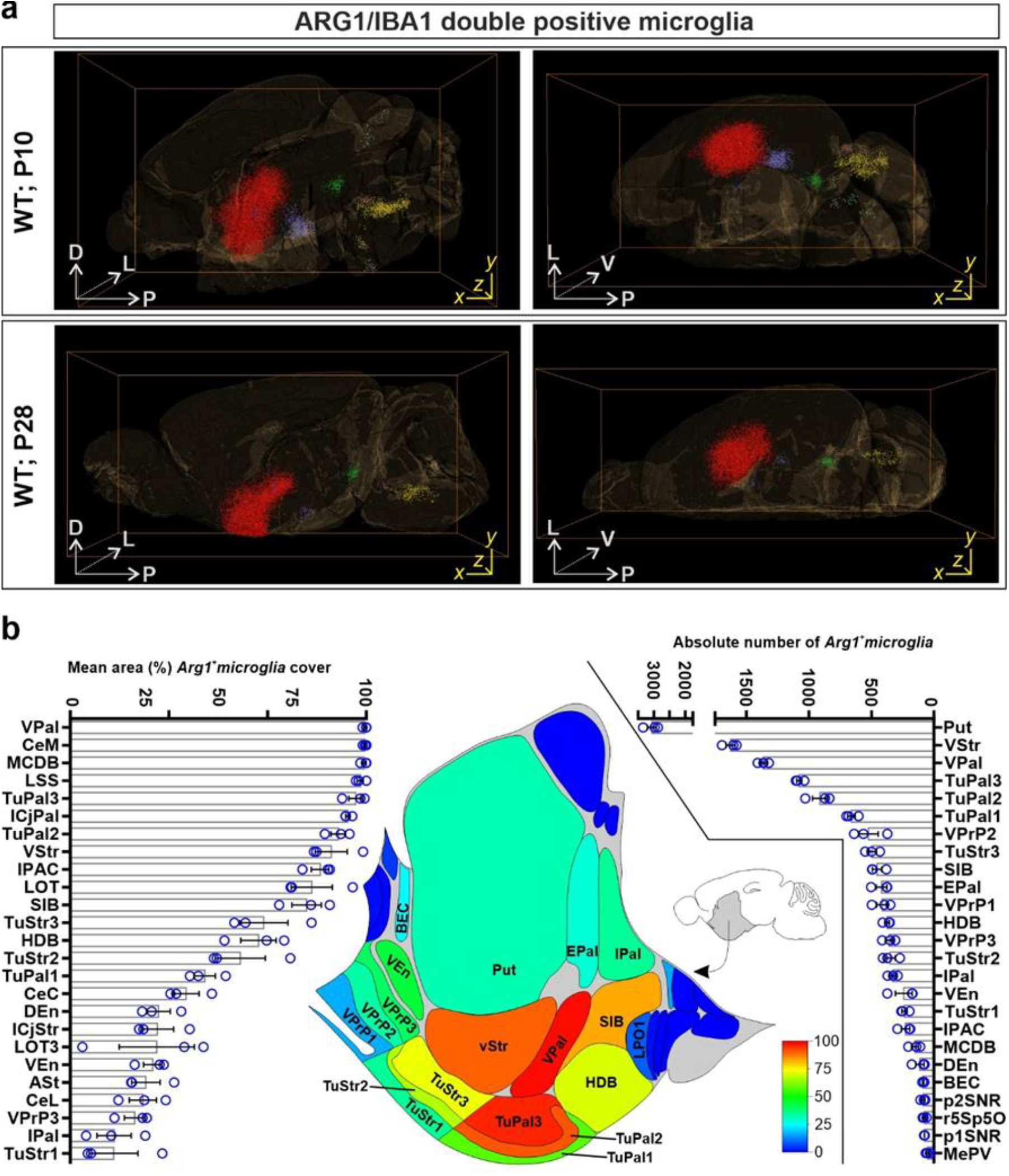
*Arg1*^*+*^ microglia have a site-specific distribution in the WT mouse brain. **a**, *Arg1*^*+*^ microglia are found in specific locations in the brain forming clusters. The largest cluster is found in the BF/vStr (red cluster). Each dot corresponds to one cell (n=1 animal). **b**, Registration of iDISCO+ to the Allen Mouse Brain map of the BF/striatum (n=3 animals). Percentage of area (pixels) that *Arg1*^*+*^ microglia occupy, accompanied by graphical illustration and absolute number of *Arg1*^*+*^ microglia in each brain area as quantified by iDISCO+. Scale bars, x=y=1000 μm; z=5580 μm (**a** (P10)) and z=5845 μm (**a** (P28)). D, dorsal; V, ventral; P, posterior; L, lateral. Data in **b** represented as mean ± s.e.m. Schematic in **b** was adapted from the Allen brain Institute Reference Atlas (http://mouse.brain-map.org).

### *Arg1*^*+*^ microglia number decreases throughout the postnatal development

While *Arg1*^*+*^ microglia were present at all investigated ages, P10, P28 and P100, their numbers varied greatly. The presence of *Arg1*^*+*^ microglia in the BF/vStr at P28 was reduced to approximately one third of the population observed at P10 (Fig. 1c), while at P100 only a residual population could be observed (Supplementary Fig. 2b-c). Of note, early postnatal life is a critical period for brain development during which brain size increases^27^ and neuronal spines and networks mature^28^, including cholinergic BF neurons^29^. Collectively these data support the existence in the unchallenged WT mouse brain of a subset of microglia, which expresses ARG1 and exhibits intriguing spatiotemporal overlap with the cholinergic system.

### *Arg1*^*+*^ microglia are morphologically indistinguishable to neighbouring “canonical” *Arg1*^*-*^ microglia

Thereafter, we wanted to explore in which terms *Arg1*^*+*^ microglia can differ from their neighbouring “canonical” counterparts, *i.e*. ARG1-non expressing. We used morphometric analysis to compare *Arg1*^*+*^ microglia to neighbouring “canonical” microglia from P10 and P28 animals, but no significant differences were observed (Supplementary Fig. 3a-b).

### *Arg1*^*+*^ microglia have an immune vigilant transcriptomic profile and do not express classical markers associated to “alternative” reactive microglia

To assess whether *Arg1*^*+*^ microglia are characterized by an individual gene expression signature, their global transcriptome was compared to that of neighbouring “canonical” microglia. For this purpose, we sorted ARG1-YFP^+^/CX3CR1^+^/CD206^-^ and ARG1-YFP^-^/CX3CR1^+^/CD206^-^ microglia from the BF/vStr of P13 YARG mice and performed high-throughput RNA sequencing (RNA-Seq) analysis. First, we validated by immunofluorescence that ARG1 and YFP protein expressions in cells are concurrent in YARG mice brain (Supplementary Fig. 2a). We also validated in CX3CR1-GFP mice, which express GFP under the control of the endogenous *Cx3cr1* locus, that a subpopulation of CX3CR1-expressing microglia expresses ARG1 (Supplementary Fig. 4a). Of note, we observed in WT brain vessels ARG1^+^/IBA1^+^ cells with amoeboid morphology, which were strongly reminiscent of perivascular macrophages^30^ (Supplementary Fig. 4b). In fact, those cells were positive for the macrophage mannose receptor (*CD206*)^30^, in contrast to the microglia which were negative for this marker and exhibited a ramified morphology. Based on the above observations, we sorted *Arg1*^+^ microglia and simultaneously collected “canonical” neighbouring microglia from an area ventral to the corpus callosum, anterior to the lateral ventricles and excluding the olfactory bulb of YARG mice (Fig. 3a). To minimize potential microglial activation, cell isolation was performed under cold conditions and using low endotoxin Percoll gradient. Subsequently, the cell populations of interest were collected by fluorescence-activated cell sorting, using negative CD206 selection (to exclude perivascular macrophage), positive selection for CX3CR1 (*i.e*. microglial marker) and ARG1-YFP expression (Fig. 3a and Supplementary Fig. 5a). Three independent biological replicates from pooled male mouse brain tissues were used for transcriptomic analysis. qPCR analysis for the sorted ARG1-YFP^+^/CX3CR1^+^/CD206^-^ and ARG1-YFP^-^/CX3CR1^+^/CD206^-^ cell populations confirmed the expression of *Cx3cr1* and *Aif1*, but most significantly *Arg1* gene expression was restricted to the ARG1-YFP^+^/CX3CR1^-^/CD206^-^ population (Supplementary Fig. 5b). RNA-Seq data revealed that the *Arg1*^*+*^ microglia possess a unique and distinct transcriptomic profile when compared to “canonical” microglia from the same brain area. Based on these genome wide gene expression analyses, 269 genes were found to be up- and 16 to be down-regulated at least 3-fold, when compared to “canonical” microglia (Fig 3b-c and Supplementary Data 1). *Arg1* was found to be one of the most differentially expressed genes between these two microglia populations. *Arg1*^*+*^ microglia also express high mRNA copy numbers for microglial homeostatic genes such as *P2ry12, Tmem119, Siglech, Gpr34, Socs3, Hexb, Olfml3* and *Fcrls*^31^, validating that these cells are indeed microglia (Fig. 3d). Noteworthy, most of these microglial homeostatic genes are expressed at lower levels when compared to “canonical” microglia from the same brain area, a feature that was recently reported for reactive^32^ and disease-associated microglia^33, 34^.

**Fig. 3.**
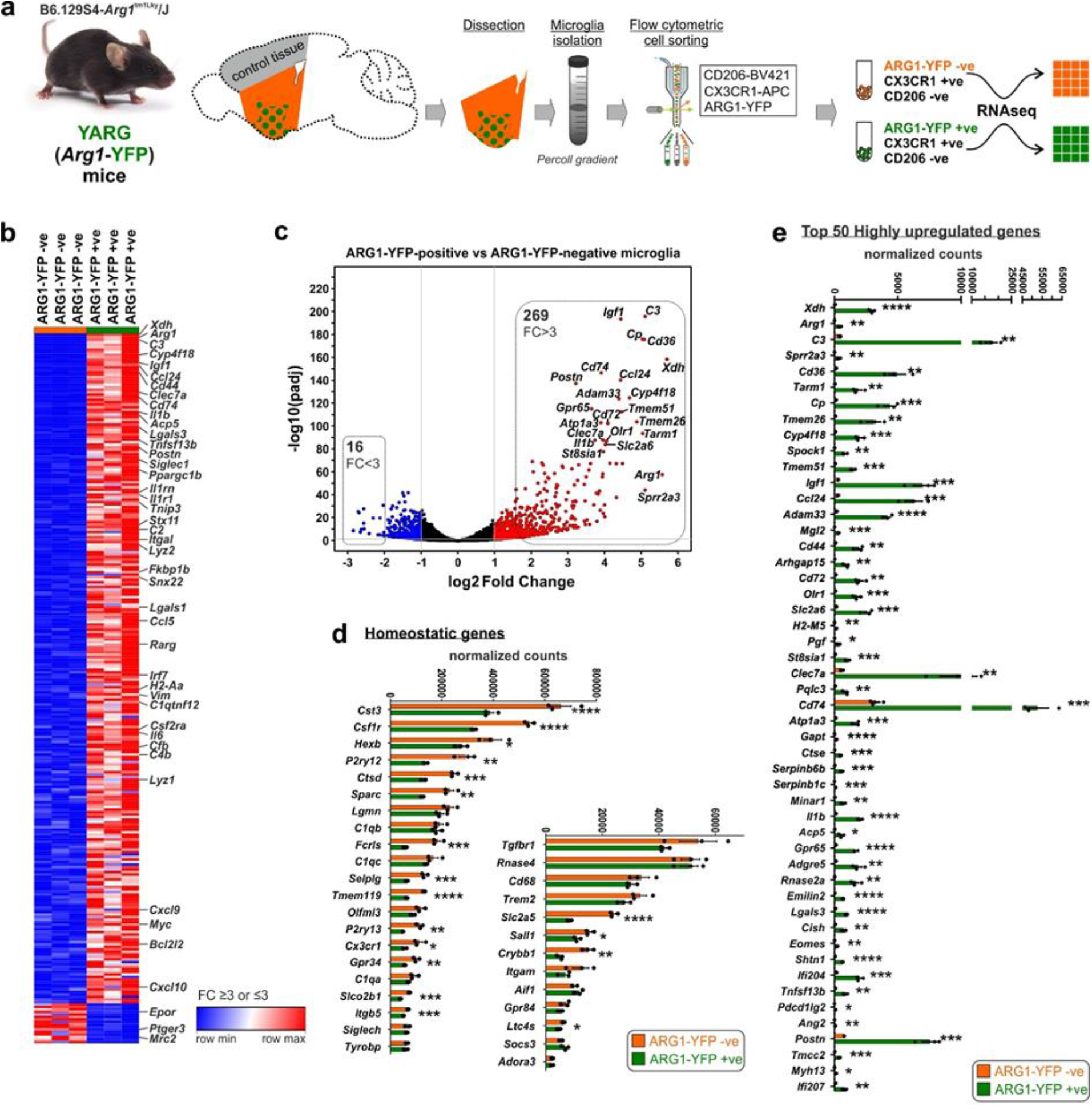
*Arg1*^*+*^ microglia transcriptome is significantly different to neighbouring “canonical” microglia. **a**, Four to five brains per biological replicate from P13 YARG male mice were dissected to cell sort *Arg1*^*+*^ microglia. Tissues were enzymatically and mechanically treated prior performing Percoll gradient centrifugation. “Canonical” and *Arg1*^*+*^ microglia were sorted by flow cytometry followed by RNA-Seq. **b-c**, Gene expression data (**b**) and volcano plot (**c**) of up- and down-regulated in *Arg1-YFP*-microglia, at least 3-fold. Note: only validated genes have been included in this list. **d**, *Arg1-YFP*^*+*^ microglia express high transcript number of homeostatic microglial genes. **e**, List of 50 most upregulated genes in *Arg1-YFP*^*+*^ microglia. Data in **d-e** are in mean ± s.e.m. Significant differences were determined by an unpaired two-tailed *t* test (\**P*< 0.05, \*\**P*≤ 0.01, \*\*\**P*≤ 0.001, \*\*\*\**P*≤ 0.0001).

Despite that *Arg1* gene expression has been traditionally associated with “alternatively” reactive microglia, RNA-Seq analysis shows that under normal conditions, *Arg1*^*+*^ microglia can neither be classified as “classically” nor as “alternatively” reactive microglia (Supplementary Fig. 5c-d). Of note, *Arg1*^*+*^ microglia are characterized by high expression of genes, including *Axl, Clec7a, Cst7, C3, Lgals3, Igf1* and *Il-1b* (reference ^31^) (Fig. 3e and Supplementary Data 1). Co-expression of Galectin-3 protein (encoded by the gene *Lgals3)* in *Arg1*^*+*^ microglia was confirmed by immunofluorescence analysis (Supplementary Fig. 5e). The visualisation of gene ontology (GO) term analysis using enrichment map showed high representation of biological processes associated with the regulation of immune and inflammatory response, immune cell communication, cell motility and chemotaxis (Fig. 4a-b). This suggests that *Arg1*^*+*^ microglia are more “immune vigilant” than “canonical” microglia. A recent high-throughput microglia single-cell transcriptome revealed several distinct microglial subsets^4^. In the context of the current investigation, the microglial “cluster 1” was found to be of particular interest. “Cluster 1” was characterized by *Arg1* gene expression, represented a small percentage (0.5%) of the whole brain microglial population and was predominantly observed at young ages. Direct comparison of the *Arg1*^*+*^ microglia and “cluster 1” genes revealed that these cells partially share a common transcriptomic signature (Supplementary Fig. 5f).

**Fig. 4.**
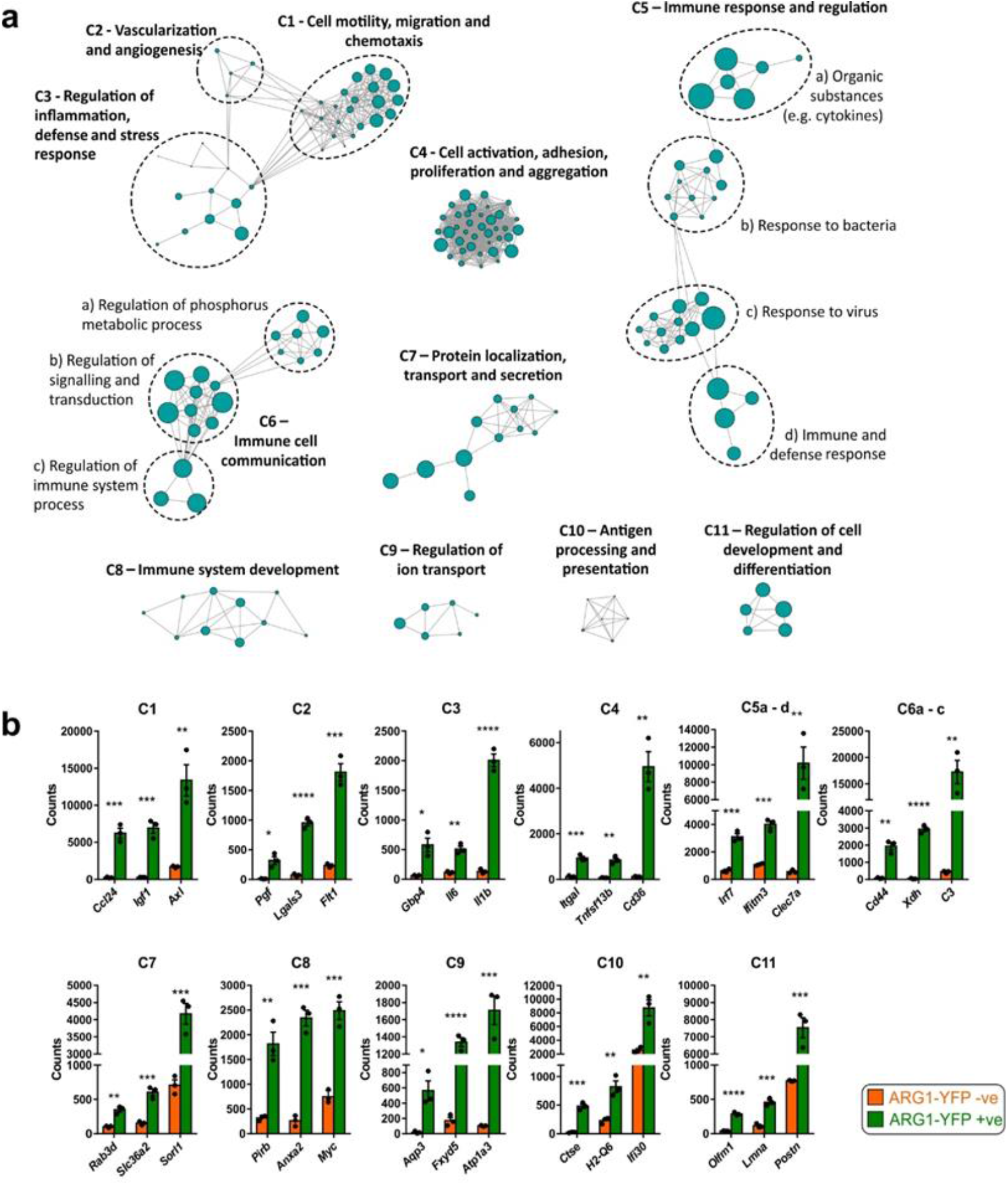
GO terms of genes highly expressed in *Arg1*^*+*^ microglia suggests greater immune vigilant and surveillance phenotype of these cells. **a**, Network graph displaying enriched GO terms of biological processes (analysed using DAVID) imported into Enrichment map, nodes = GO terms, edges = relation between GO terms above the Jaccard coefficient of 0.5, cluster size ≥5. **b**, Representation of example genes contained in distinct enrichment map clusters. Data in **b** are in mean ± s.e.m. Significant differences were determined by an unpaired two-tailed *t* test (\**P*< 0.05, \*\**P*≤ 0.01, \*\*\**P*≤ 0.001, \*\*\*\**P*≤ 0.0001).

### *Arg1* microglial conditional knockout results in impaired cognitive function in female mice

Given the unique spatiotemporal distribution and significantly different transcriptome of *Arg1*^*+*^ microglia when compared to neighbouring “canonical” microglia, we sought to investigate if they also have a distinct functional specialization. We specifically knocked out *Arg1* in microglia by crossing *Cx3cr1*^*CreER*^ with *Arg1*^*flox/flox*^ mice and induced recombination by multiple tamoxifen injections from P1 (Fig. 5a-b). Using immunohistochemistry, we confirmed that ARG1 was knocked down efficiently (Supplementary Fig. 6). Because of the topographic organisation of *Arg1*^*+*^ microglia proximal to the cholinergic nucleus of BF, a major nucleus for cognition^25^, we prompted to investigate if cognition in conditional *Arg1*-knockout animals (*Arg1*^*flox/flox*^.*Cx3cr1*^*CreER+/-*^; thereafter referred as *Arg1*^*cKO*^) compared to controls (*Arg1*^*flox/flox*^*CX3CR1*^*CreER-/-*^ ; thereafter referred as *Arg1*^*Control*^) is affected. Two- to three-month-old *Arg1*^*cKO*^ female and male mice did not display motor-coordination dysfunction in comparison to *Arg1*^*Control*^ (Fig. 5c-e and Supplementary Fig. 7a-c). To study hippocampus-dependent spatial memory, we used the novel arm discrimination (spatial recognition memory) paradigm in Y maze. This test is based on the inherent preference of mice to explore a novel environment more than a familiar one. The spontaneous alternation triplets percentage was similar in female (Fig. 5f) and male experimental groups (Supplementary Fig. 7d), indicating that the working memory seems to be unaffected when arginase is depleted in microglia. Interestingly, *Arg1*^*cKO*^ female mice showed a statistically significant decrease in the percentage of times they visited the new arm, when compared to *Arg1*^*Control*^ (Fig. 5g), meaning that the absence of the *Arg1*^+^ microglia impair cognitive function in the mouse model. To gain further insights, we performed object recognition memory (ORM) test. *Arg1*^*cKO*^ female mice had a reduced preference for a new object compared to a familiar one 24 hours after the training session, therefore exhibiting impairment in long term memory (LTM) acquisition (Fig. 5h). No differences were found in short term memory (STM) index (Fig. 5h) measured 1 hour after training protocol, indicating once more that the working memory was unaffected in the experimental groups. In contrast, we were not able to detect any behavioural phenotype in *Arg1*^*cKO*^ male mice (Supplementary Fig. 7e). Whether the observed behaviour difference between male and female mice upon microglial *Arg1*-knockout could be linked to the reported regulation of arginase enzymes by steroid hormones^35-37^ requires further investigation. Yet, similar gender differences have been reported when *Arg1* is depleted in peripheral myeloid cells^38^.

**Fig. 5.**
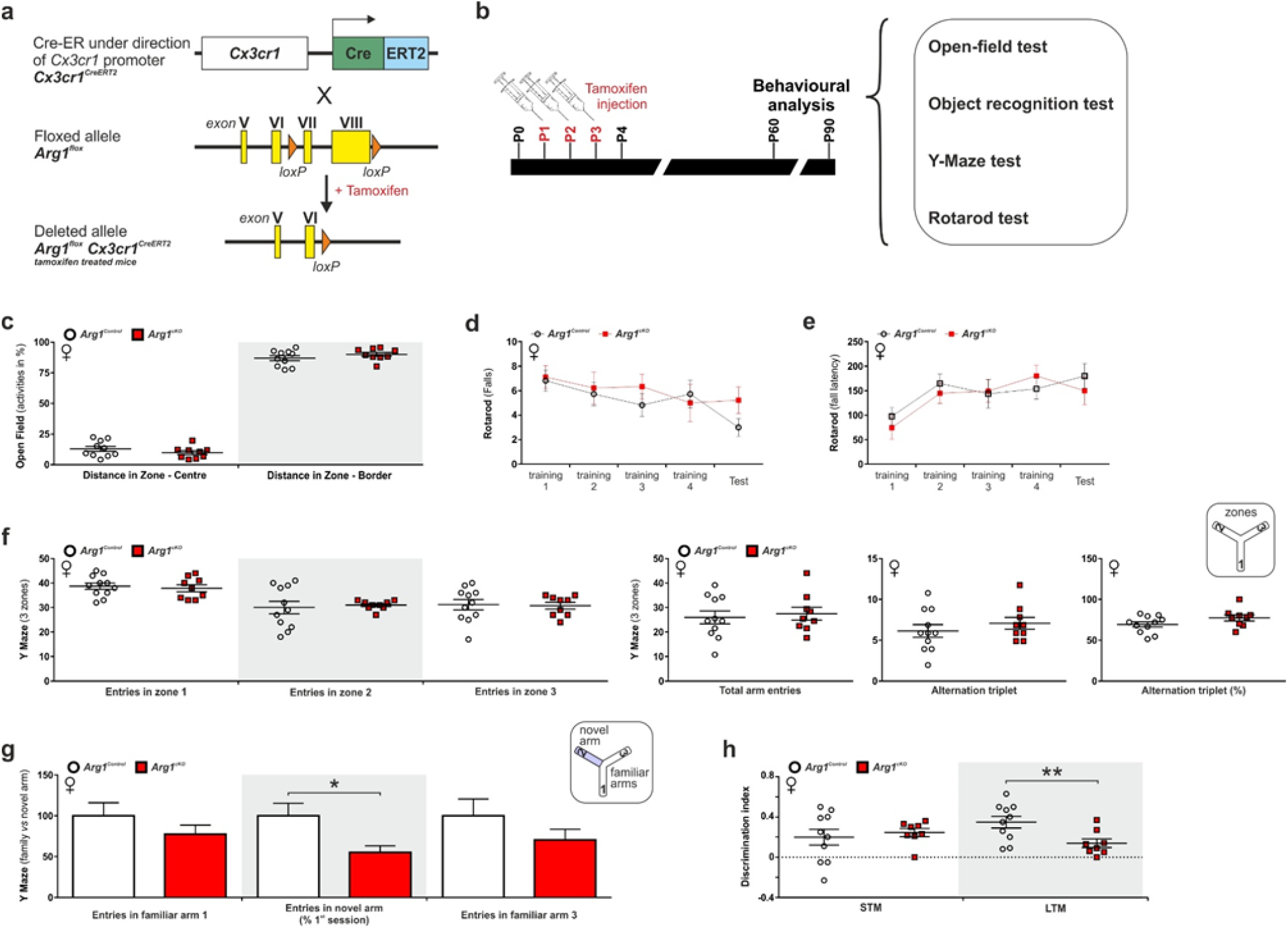
*Arg1* microglial conditional knockout leads to an impaired cognition phenotype in female mice. **a-b**, Strategy for *Arg1* conditional knockout and subsequent behavioural studies. **c-h**, *Arg1*^*cKO*^female animals and controls were assessed for motoric (**c-e**) and memory (**f-h**) phenotypes. *Arg1*^*Control*^ n=10 (for **c-e**,**h**), n=11 (for **f-g**) and *Arg1*^*cKO*^ n=9 (for **c-g**), n=8 for **h**. Data are in mean ± s.e.m. Significant differences were determined by an unpaired two-tailed *t* test (\**P*< 0.05).

### *Arg1*^*+*^ microglia are frequently in contact with neurons and contain cellular inclusions

Ultrastructural analysis of *Arg1*^*+*^ microglial cell bodies in BF/vStr of female mice revealed their frequent contacts with neurons, excitatory synapses and myelinated axons, implying an active involvement with neural development (Fig. 6a-c). In particular, half of the *Arg1*^*+*^ microglia were found in close apposition to at least one neuronal cell body, with their cell body occupying satellite positions. Interestingly, subsets of perineuronal satellite microglia have been reported earlier on, and proposed to either promote plasticity or to provide trophic support to neurons^39, 40^. Each microglial cell body also made one or more close appositions with synaptic clefts, where synaptic transmission occurs between pre-synaptic terminals and post-synaptic spines. Intrinsic features of the *Arg1*^*+*^ microglia also comprise cellular inclusions containing debris, indicating that these cells are phagocytic (Fig. 6a,c). Proximity of *Arg1*^*+*^ microglia to cholinergic neurons was verified by immunohistochemistry (Fig. 6d).

**Fig. 6.**
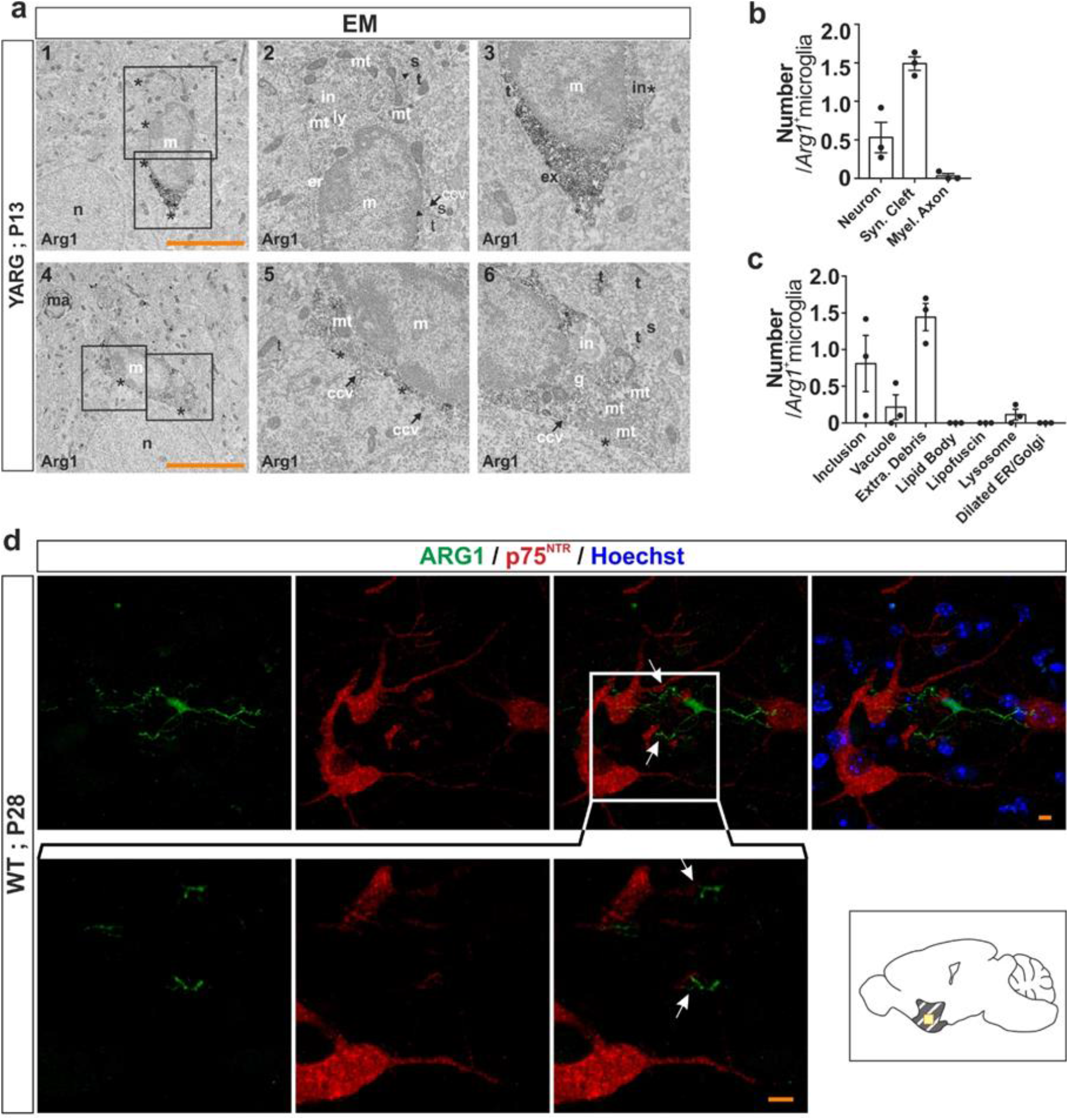
*Arg1*^*+*^ microglia are in frequent apposition to neuronal cell bodies and contain cellular inclusions. **a**, Transmission electron microscopy of *Arg1*^*+*^ microglia of P13 YARG female brains at BF/vStr. **b-c**, Quantitative analysis of intracellular relationships (**b**) and intracellular features (**c**) (n=3 female animals, 33 cells in total). **d**, In WT, P28 BF/vStr, *Arg1*^*+*^ microglia are in close proximity to cholinergic neurons, as these were detected with antibody against p75^NTR^. Scale bars, 5 μm, z=12 and 2 μm (for (**d**) upper and lower panel, respectively). Yellow squares indicate location of the corresponding images on their left. Arrowhead, synaptic cleft; Asterisk, plasma membrane; ccv, clathrin-coated vesicle; d, dendritic spine; er, endoplasmic reticulum; ex, extracellular space pocket; g, Golgi apparatus; in, phagocytic inclusion; ly, lysosome; m, microglial cell body; ma, myelinated axon; mt, mitochondrion; n, neuronal cell body; t, axon terminal (for **a**). Data are in mean ± s.e.m.

### *Arg1* microglial conditional knockout female mice have less mature hippocampal spines

To examine the cellular mechanism of the cognition phenotype, we analysed dendritic spines of CA1 and dentate gyrus (DG) hippocampal neurons in P60 female mice, respectively (Fig. 7). Hippocampus is a structure important for cognitive functions which is innervated by BF cholinergic neurons^25^, while spine plasticity has a major role in cognitive functions^28, 41, 42^. Immature spines are thinner and form less stable synaptic contacts compared to mature mushroom shaped spines ^28, 42^ (Fig. 7e), while spine density provides an estimate of synapse density^43^. More specifically, we analysed segments of secondary dendrites of pyramidal neurons located at the level of *stratum radiatum* (Fig. 7a) and found that these spines in *Arg1*^*cKO*^ female animals are significantly narrower and longer when compared to *Arg1*^*Control*^ (Fig. 7c). Morphological analysis of the pyramidal spines showed that P60 *Arg1*^*cKO*^ female have an increased proportion of immature spines (filopodia and long thin spines) and a decreased proportion of mature spines (mushroom and branched) (Fig. 7e-f). Analysis of the granule ce**l** s’ dendritic spines of the outer third of the suprapyramidal blade (Fig. 7b) indicated that spine density was higher in P60 *Arg1*^*cKO*^ females (Fig. 7d). Similarly, to the pyramidal spines, secondary granule cell dendrites in P60 *Arg1*^*cKO*^ females were significantly narrower (Fig. 7d) and had a higher percentage of immature and lower percentage of mature spines (Fig. 7e,g). Collectively these data indicate that the impaired cognitive development observed in the *Arg1*^*cKO*^ female mice is related to disruption of maturation of hippocampal dendritic spines.

**Fig. 7.**
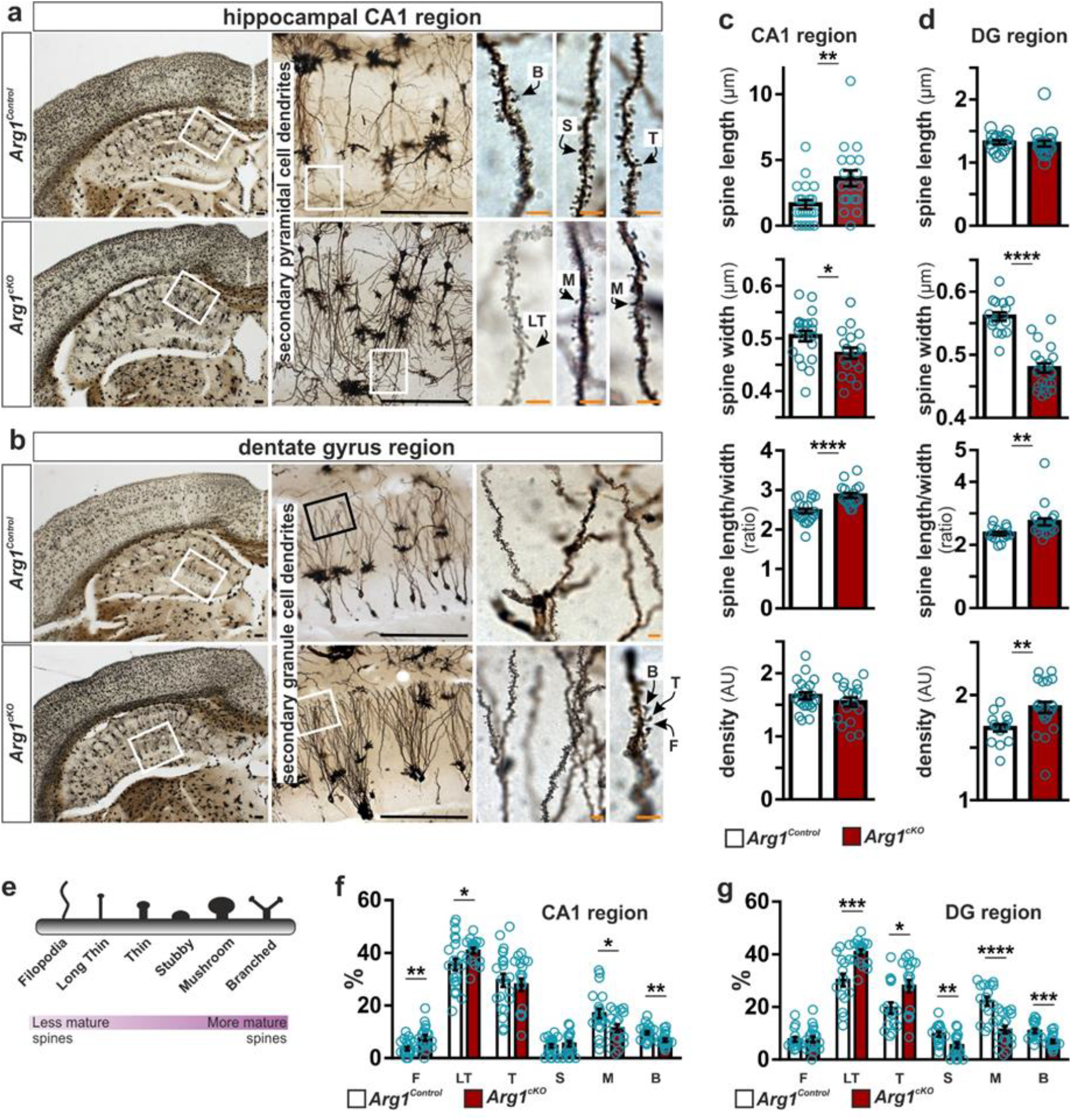
*Arg1* microglial conditional knockout affects dendrite maturation in the hippocampus. **a-b**, Coronal Golgi-Cox-stained sections of P60 female *Arg1*^*cKO*^ brains and controls. Hippocampal CA1 and the DG regions from which secondary pyramidal and secondary granule cell dendrites, respectively, were used for spine counts. **e**, Six main spines categories according their morphological characteristics are: filopodia, long thin, thin, stubby, mushroom, and branched. (**c-g**) Graphical representation of differences between spines in hippocampal CA1 (**c**,**f**) and DG (**d**,**g**). F, filopodia; LT, long thin spines; T, thin spines; M, mushroom spines; S, stubby spines (for **f**,**g**). Black scale bars, 200 μm, orange scale bars, 5 μm. Data are in mean ± s.e.m. Significant differences were determined by an unpaired two-tailed *t* test (\**P*< 0.05, ** *P*≤ 0.01, \*\*\**P*≤ 0.001, **** *P*≤ 0.0001).

## Discussion

During development, microglia have pivotal roles in all aspects of neuronal and glial physiology, including control of neurogenesis and oligodendrogenesis, refinement of neuronal wiring, synaptic pruning, maturation, and plasticity^44^. Given the range of microglial functions, it is not surprising that conditions resulting in microglial impairment during development lead to a wide spectrum of neurodegenerative and neuropsychiatric disorders^44, 45^. Compiling data show that microglia are not a homogenous population that responds stereotypically to extrinsic stimuli, but rather a heterogeneous cell type that exhibits transcriptomic profile^1-5^ and ultrastructural^46^ differences to name a few, a diversity that is especially pronounced during early postnatal development (reviewed in ^6^). Here we report a microglial subtype morphologically indistinguishable from neighbouring “canonical” microglia, but exhibiting a significantly different transcriptome, dynamic spatiotemporal localization and functional specialization (Fig. 8). We show that the enzyme ARG1 is highly expressed in this microglial subpopulation and is essential for proper brain development. We provide here compelling evidence for a critical role of *Arg1*^+^ microglia in shaping neuronal circuits involved in cognition. Supporting this, microglial *Arg1*-knockout results in impaired neuronal plasticity and cognitive deficits in mice. Previous studies have shown that whole body *Arg1-* knockout mice are postnatally lethal and exhibit neurotoxicity^47^. In humans, ARG1-deficiency is a rare autosomal disease, more frequently diagnosed in infant females than males, in which the phenotype is less severe in patients surviving to adulthood^47, 48^. As in mice, ARG1-deficient patients show neurological problems evident by the progressive neurological and cognitive impairment leading to various degrees of intellectual disability^47^. Most interestingly, the vast majority of the patients whose peripheral symptoms can be managed through diet or drug therapy, still continue to suffer from cognitive deficits^47^. Whether the human pathology is linked specifically to ARG1-deficiency in microglia needs to be investigated. The identification of microglial subtypes with intrinsic differences will elucidate new prospects in understanding brain physiology during homeostasis and pathology. Moreover, microglial subtypes can be *de facto* new targets for development of therapeutic strategies.

**Fig. 8.**
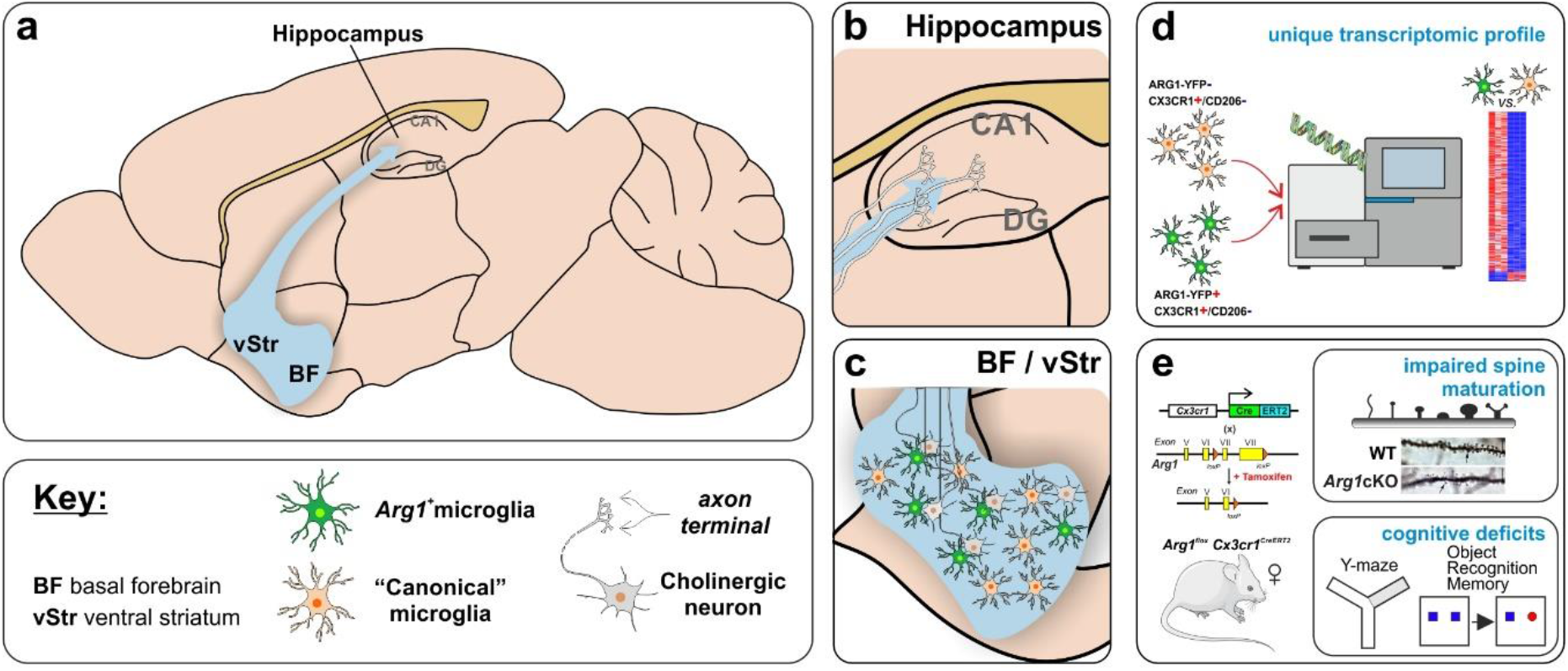
Summary of the main findings. **a-c**, The BF is a brain area rich in cholinergic somas which project to the hippocampus (**b**). *Arg1*^*+*^ microglia are found in cholinergic-soma-rich areas and most predominantly in the BF/vStr (**c**). **d**, *Arg1*^*+*^ microglia have a unique transcriptomic profile compared to neighbouring “canonical” microglia. **e**, *Arg1*^*cKO*^ female have impaired spine maturation in the hippocampus and exhibit cognitive deficits.

Our RNA-Seq analysis of “canonical” and *Arg1*^+^ microglia showed robust expression of typical homeostatic genes including, among others, *Tmem119, Cx3cr1, Hexb, P2ry12, Csf1r* and *Siglech*^32-34, 49, 50^. A recent study has demonstrated that postnatal microglia are enriched for Cst3 (encoding cystatin C) and Sparc (encoding secreted protein acidic and rich in cysteine)^1^. Both genes appear as two of the most upregulated ones in our RNA-Seq analysis, which was performed at P13, a clear indication that our cell sorting strategy was quite specific for isolating microglia. Recent massive transcriptomic analysis of microglia have identified several microglia subpopulations during early postnatal development^1, 3, 4, 18^. The most distinctive brain microglia subtype during postnatal development was originally identified by their high expression of CD11c and IGF1 and associated to the corpus callosum and cerebellar white matter^18^. Subsequent studies using massive transcriptomic analysis of microglia at the single cell level confirmed the existence of this microglia subpopulation and further defined as tract-associated microglia^4^ or proliferative-region-associated microglia^3^. Intriguingly, this microglia subtype was characterized by downregulation of homeostatic microglia markers along with upregulation of several genes typically found under disease conditions (disease-associated microglia)^3, 4, 33, 34^, thus sharing some molecular features with the *Arg1*^*+*^ microglia subtype. However, tract-associated microglia fail to upregulate *Arg1* (references ^3, 4^), which together with the high enrichment of this microglia subpopulation in the axon tracts and their amoeboid morphology full contrast with that shown in the *Arg1*^*+*^ microglia subtype^3, 4^. We may wonder how this microglia subtype has escaped its identification from most transcriptomic studies performed during postnatal development. Strikingly, Stevens and colleagues recently identified a small microglia cluster (“cluster 1” - the smallest identified cluster comprising about 0.5% of total microglia), which displays strong upregulation of *Arg1* (reference ^4^). Although there are some differences when comparing directly the results from Hammond *et al*. “cluster 1” single cell sequencing data with the *Arg1*^*+*^ microglia RNA-Seq data described here (Supplementary Fig. 5f), these could be due to a number of factors including age of animals used (P4/P5 *vs* P13), cell isolation protocol and sequencing technology. Nevertheless, both “cluster 1” and *Arg1*^*+*^ microglia display a strong upregulation of *Arg1, Cd74, Ccl24, Gatm, Clec4a3, Dmrtb1, Fcgr2b* and *Fcgr4* and are more prominent during early postnatal stages^4^, thus validating independently the *Arg1*^*+*^ microglia subtype. In addition, “cluster 1” size shows sex-specific variation being enriched in female mice^4^. This observation is significant as it provides strong support to the expanding literature about sex differences in microglia^19-22^, while it can partially explain the differences in the behavioral studies we observed in this study. Taken together, we can conclude that the *Arg1*^*+*^ microglia is a developmentally regulated subtype that displays a distinctive molecular signature and has a specific function in brain shaping.

## Methods

### Animals

All animal experimental protocols in the present study were in accordance to the respective national, federal and institutional regulations, *i.e*. the Guidelines of the European Union Council, following Swedish regulations for the use of laboratory animals and approved by the Regional Animal Research Ethical Board, Stockholm, Sweden (Ethical permits N248/13) the Spanish regulations (BOE 34/11370–421, 2013) and in conformity with the Canada Council on Animal Care guidelines. C57BL/6J mice (WT, Charles River, Sulzfeld, Germany), *YARG* (The Jackson Laboratory, stock # 015857), *CX3CR1-GFP* (The Jackson Laboratory, stock # 005582), *Arg1*^*flox/flox*^ (The Jackson Laboratory, stock # 008817) and *CX3CR1*^*CreER*^ (The Jackson Laboratory, stock # 021160) mice were used and maintained under a 12-h light/dark cycle at 22–25°C with access to food and water *ad libitum*.

### Generation of microglia specific Arg1-deficient mice (Arg1^cKO^)

*Arg1*^*flox/flox*^ (The Jackson Laboratory, stock # 008817) mice with the *Arg1* allele bearing *loxP* sites flanking exons 7 and 8 were crossed with *CX3CR1*^*CreER*^ (The Jackson Laboratory, stock # 021160) to generate *Arg1*^*flox/flox*^*CX3CR1*^*CreER+/-*^ (*Arg1*^*cKO*^) and *Arg1*^*flox/flox*^*CX3CR1*^*CreER-/-*^ (control mice; *Arg1*^*Control*^) (Fig. 5a). Deletion was induced upon tamoxifen daily treatment for two or three consecutive days starting at P1 (Fig. 5b). All mice (Cre+ and Cre-) were injected with tamoxifen at the following doses: 250 μg/pup. Genotyping of the mice was done by PCR analyses of finger DNA using the following primers for the *Arg1* floxed allele: TGCGAGTTCATGACTAAGGTT as forward primer and AAAGCTCAGGTGAATCGG as reverse one. To detect the presence of CRE we used three specific primers which amplified both WT (AGGATGTTGACTTCCGAGTTG) and CRE band (CGGTTATTCAACTTGCACCA) and the common primer was AAGACTCACGTGGACCTGCT.

### Tissue preparation, immunohistochemistry and confocal laser microscopy

Female animals (unless otherwise specified) were deeply anesthetized with sodium pentobarbital, transcardially perfused with 0.9% sodium chloride, and then fixed with 4% paraformaldehyde in 0.1M phosphate buffer (pH 7.4). Brains were collected and postfixed in the same fixative for 24 h, then transferred to 30% sucrose in 0.1 M phosphate buffer for cryoprotection, and left for a minimum of 3 days. Brains were cryosectioned sagittally using a sliding microtome (Leica SM2000R) into 25 µm free floating sections and stored as 1:12 series at 4° C in tubes containing cryoprotection solution (25% glycerin, 25% ethylene glycol in 0.1M phosphate buffer) for further histological analysis. When necessary, antigen retrieval was performed by heat induced antigen retrieval (DAKO, S1699), followed by 3×10 min PBS, 1×20 min PBS/0,3 % Triton-X and 1 hour blocking in PBS/0,3 % Triton/10% donkey serum, all steps at room temperature (RT). Stainings were performed consecutively. Primary antibodies used in this study: ARG1 (1:400, rabbit, Abcam, ab91279), ARG1 (1:100, goat, SantaCruz, sc-18354; 1:100, mouse, sc-271430 in Supplementary Fig. 4b), CD206 (1:100, goat, R&D, AF2535), GALECTIN-3 (1:400, goat, R&D, AF1197), GFP (1:50, goat, Abcam, ab6673), IΒΑ1 (1:250, goat, Abcam, ab5076), IΒΑ1 (1:250, rabbit, Wako, 01919741) and p75^NTR^ (1:1000, rabbit, G3231, Promega). Sections were washed 3×20 min in PBS prior to incubation with secondary antibodies (Alexa Fluor) for 1 hour at RT. Finally, sections were washed 3×5 min win PBS, counterstained with 0.1 μg/ml Hoechst, further washed 3×5 min in PBS and mounted in Fluoromount-G (SouthernBiotech, 0100-01). Images were acquired with a Zeiss LSM700 confocal laser scanning microscope and assembly was performed with ImageJ/Fiji, Adobe Photoshop and CorelDraw.

### Manual cell number quantification

The number of *Arg1*^*+*^ microglia, based on ARG1 expression, from four consecutive matched sections within the 1:12 series was counted and added to give the relative number. Four animals per age group (Fig. 1c) and two animals per genotype (Supplementary Fig. 6b) were used. For the percentage of *Arg1*^*+*^ microglia quantification, the area *Arg1*^*+*^ microglia cover was marked and the percentage of *Arg1*^*+*^ microglia versus IBA1^+^ microglia was quantified. Researchers performing the quantifications were blinded to the samples.

### iDISCO+ immunostaining and tissue clearing

iDISCO+ immunostaining and tissue clearing was performed as previously published^24, 51^. P10 and P28 WT male mice brains were perfused using 4% paraformaldehyde and fixed overnight at 4 °C. After washing the brains with PBS, the samples were cut into half hemispheres and treated with a series of methanol solution (20, 40, 60, 80, 100, 100%) and stored in -20 °C until use. The samples were treated with 5% hydroxyperoxide in methanol at 4 °C overnight, then treated with reverse concentration of methanol solution (80, 60, 40, 20%), and washed wish iDISCO washing buffer (PBS with 0.2% Tween-20 and 10 µg/ml Heparin). After permeabilization (PBS with 0.2% TritonX-100, 0.3 M Glycine, 20% Dimethyl Sulfoxide(DMSO)) and blocking (PBS with 0.2% TritonX-100 with 6% Donkey serum, 10% DMSO) the samples for 2 days each, they were incubated with primary antibodies, namely ARG1 (1:50, goat, SantaCruz, sc-18354) and IBA1 (1:200, rabbit, Wako, 01919741) for 5-7 days at 37 °C with gentle rotation. After washing the samples with iDISCO washing buffer for one day, they were incubated with secondary antibodies for 5-7 days at 37 °C with gentle rotation. The samples were washed with iDISCO wash buffer for one day with gentle rotation at 37 °C. The immunostained hemispheres were treated with an up-series of methanol solution (20, 40, 60, 80, 100, 100%) at RT. The samples were incubated in 33% methanol / 66%DCM (Dichloromethane, Sigma, 270997) for 3 hours and 100% DCM for 15 min twice, then transferred in DBE (Benzyl ether, Sigma, 108014). After one day incubation of DBE, the DBE solution was changed to a fresh one and incubated for a day before imaging.

### iDISCO+ imaging acquisition

Cleared brain images were acquired by the COLM microscope^52^. The hemisphere was placed in quartz cuvette (Starna Scientific) filled with DBE and fixed position using silicone blocks (Sylgard 184, Dow Corning). Sagittal brain images were acquired. Immunostaining signal was acquired by 647 or 561nm channel, and autofluorescence images were acquired using 488nm or 405nm laser channel for image registration. Original image resolution is 0.585, 0.585, 5 μm in x, y and z axis.

### iDISCO+ image analysis

Original tiff file (16bit) was downsampled to 5μm isotropic resolution and converted into 8bit by custom MATLAB script and Fiji^53^. Each z-stack image was stitched by TeraStitcher^54^. The stitched images were processed by a series of processing filters (unsharp mask, background subtraction, integral filter and fast Fourier transform filter) in ImageJ and Amira 3D software (ThermoFisher Scientific). ARG1^+^ and IBA1^+^ cells were recognized by signal intensity and morphology. After removing blood vessels manually in Amira segmentation editor, ARG1^+^ and IBA1^+^ cells were segmented by global threshold segmentation. Segmented images were labelled, counted, and converted into points cloud by Amira. Movies were generated using Amira.

### Image registration

Image registration was performed using Elastix toolbox^55, 56^ and MelastiX MATLAB wrapper (https://github.com/raacampbell/matlab_elastix). Four different methods of image registration (Rigid, Similarity, affine, B-Spline) were sequentially performed to register sample data. First, we acquired reference brain images (P28 and P10) stained with Neurotrace 640/660 (N21483, ThermoFisher Scientific). Next, we transformed the reference brain image to Allen brain atlas (Developing mouse, P56 and P14, June 2013 v.2) using Neurotrace 640/660 channel and obtained autofluorescent image registered to Allen brain atlas (AutoF Allen brain atlas). Lastly, immunostained brain images were registered to the AutoF Allen brain atlas using autofluorescence channel of the immunostained brain.

### Morphometric analysis

25 μm free floating sections were stained with IBA1 (1:250, rabbit, Wako, 01919741) and ARG1 (1:100, goat, SantaCruz, sc-18354) antibodies and individual microglia from BF/vStr were selected for 3D reconstruction and scanned with Zeiss LSM700 with the following parameters: 1024 × 1024 pixel resolution, 0.5 μm step, x63 objective. Ce**l** s to be imaged needed to have their nucleus in the centre of the z-plane. For quantitative microglia morphology analysis, confocal Z-stacks were automatically reconstructed using a self-customized python-based script^57^. Reconstructions were visually checked using the ImageJ plugin “simple neurite tracer”. Each ce**l** was individually extracted, and the number of branches, path length, branch order and processes volume were quantified using the open source software L-measure. 4 mice per developmental stage were examined. At least 15 (P10) and 32 (P28) microglia of each phenotype were analysed. Quantification was performed blinded.

### Immunoperoxidase staining for electron microscopy

*YARG* mice were housed under a 12 h light-dark cycle at 22–25°C with free access to food and water. Three P13 female pups were anesthetised with a mixture of Ketamine-xylazine (80 and 10 mg/kg, i.p.) and transcardially perfused with phosphate-buffered saline (PBS; 50 mM, pH7.4) followed by 3.5% acrolein and 4% paraformaldehyde. Fifty-micrometer-thick coronal sections of the brains were cut in PBS using a Leica VT1000S vibratome (Leica Biosystems, Concord, ON, Canada) and stored at -20°C in cryoprotectant until further processing ^58^.Brain sections selected in the basal forebrain were rinsed in PBS, then incubated in 0.1 M citrate buffer for 40 min at 70 °C for antigen retrieval. They were quenched with 0.3% hydrogen peroxide (H_2_O_2_) for 5 min followed by 0.1% sodium borohydride (NaBH_4_) for 30 min. Afterwards, sections were washed three times in PBS and blocked in 10% donkey serum with 0.03% Triton X-100 in PBS for 1 h at room temperature and then were incubated overnight with a primary anti-ARG1 antibody (1:100, goat, SantaCruz, sc-18354). The next day, sections were rinsed three times in TBS and incubated for 1 h with secondary antibody conjugated to biotin (1:200, Jackson ImmunoResearch Laboratories) and for 1 h with Vectastain® Avidin-Biotin Complex Staining kit (Vector Laboratories). Sections were developed in a Tris buffer solution (TBS; 50 mM, pH 7.4) containing 0.05% diaminobenzidine and 0.015% H_2_O_2_ and then rinsed with PBS. The sections were post-fixed with 1% osmium tetroxide, dehydrated using sequential alcohol baths followed by propylene oxide. Sections were embedded in Durcupan resin (Sigma-Aldrich) between ACLAR sheets at 55°C for 3 days, as described previously^59^. Ultrathin sections were generated at ∼65 nm using a Leica UC7 ultramicrotome. Microglia immunoreactive for Arg1 were imaged at 80 kV using a transmission electron microscope (FEI Tecnai Spirit G2) and photographed at a magnification of 6800X with a Hamamatsu ORCA-HR digital camera (10 MP). Due to the limited penetration of antibodies in immunoEM, we could not be affirmative when cells appeared Arg1-negative that they are “canonical” microglia; hence we did not proceed to the comparison between the two subtypes.

### Ultrastructural analysis

Only microglia stained with ARG1 and observed within the brain parenchyma were included in the analysis. The cell bodies of 10-12 microglia per mouse were analysed quantitatively. Profiles of neurons, synaptic elements, microglia, and myelinated axons were identified according to well-established criteria. The parameters examined were the prevalence of lipofuscin granules, lipid bodies, ER/Golgi dilation, lysosomes, phagocytic inclusions, vacuoles, and association with extracellular debris. In addition, the intercellular relationships of Arg1 microglia with neuronal cell bodies, excitatory synapses and myelinated axons were quantified. Neurons were identified according to their large and round nucleus, the presence of an electron dense nucleolus, the large dendrites extending from their cell body and the absence of bundles of gliofilaments.

### Golgi-Cox staining

Segments of secondary dendrites of pyramidal neurons located at the level of the *stratum radiatum* (CA1) and the granule ce**l** s’ dendritic spines of the outer third of the suprapyramidal blade (DG) were considered for quantitative analysis. Dendrite segments from 15 to 20 µm in length completely filled by the mercuric reaction of the Golgi-Cox method were used. The criteria used to categorize and quantify the dendritic spines on CA1 pyramidal neurons were previously described^42^ (Fig. 7e). 3 females for each genotype were analysed *i.e. Arg1*^*cKO*^ (P60) and *Arg1*^*Control*^ (P60). Number of secondary dendrite’s segments in CA1 counted is: 21 for *Arg1*^*Control*^ (P60) and 20 for *Arg1*^*cKO*^ (P60). Number of dendrite’s segments in DG counted is: 17 for *Arg1*^*Control*^ (P60), 22 for *Arg1*^*cKO*^ (P60).

### Microglia isolation

Brains from 4-5 P13 *YARG* male mice were perfused with 10 ml cold PBS. The olfactory bulb was removed and the two hemispheres were separated. Next, the median plane of each hemisphere was placed facing upwards and a cut was made posterior to the lateral ventricle and the tissue ventral to corpus callosum (including the cerebral nucleus) was dissected out. The remaining tissue (corpus callosum and cortex) was collected and used as control tissue. Tissues were pooled, minced with scalpel, further dissociated with enzymatic solution (0,01% Papain, Sigma, 10108014001; 0,1% dispase II, Sigma, D4693; 0,005% DNAseI, Sigma, 10104159001; 12,4 mM MgSO_4_ in HBSS) for 20 min at 37 °C and filtered through 70 µm nylon mesh (Miltenyi, 130098462). Microglial fraction was enriched by 20% Percoll density gradient (Percoll PLUS, GE Healthcare, 17544502) at 4 °C.

### Fluorescence-Activated Cell sorting

Microglial population was initially gated based on size and granularity, followed by gating for singlets, negative selection with CD206-BV421 (Biolegend, 141717) to exclude macrophages and positive selection with CX3CR1-APC (R&D, FAB5825A). Finally, microglia were sorted to *Arg1*^*+*^ and *Arg1*^*-*^ based on YFP expression. Sorting was performed on FACSAria III Cell Sorter system and analysed using FACSDiva™ software (BD Biosciences). Ce**l** populations (ARG1-YFP^+^/CX3CR1^+^/CD206^-^, ARG1-YFP^-^/CX3CR1^+^/CD206^-^ and CD206^+^) were collected directly in Qiazol reagent (Qiagen) followed by RNA extraction (using the RNeasy micro kit, Qiagen). RNA quality and concentration was assessed by the sequencing facility using Bioanalyser before library preparations and sequencing.

### RNA isolation and qPCR

RNA was isolated using commercial kits (RNeasy micro, #217084, Qiagen). cDNA was synthesized using Oligo dT, dNTPs and Superscript III (Invitrogen). qPCR was performed using the StepOne plus instrument (Applied Biosystems) using the SYBR™ Green master mix (life technologies) and predesigned primers (KiCqStart® Primers, Sigma). Relative gene expression levels were normalized to β-actin in each sample with the ΔΔCT method.

### Library preparation and RNA-Seq

2 ng of total RNA was used to generate cDNA for the samples using the Smart seq V4 Clontech reagents. Libraries were prepared with the Illumina Nextera XT protocol. The library pools underwent cluster generation on an Illumina cBot and the clustered flow cell was sequencing on the IlluminaHiSeq 2000 instrument to generate 50 bp SR. The raw sequencing reads were processed through CASAVA for FASTQ conversion and demultiplexing.

### RNA-Seq data and computational analysis

Basecalling and demultiplexing was performed using Illumina bcl2fastq v2.20.0 software. Reads were aligned to Ensembl GRCm38/mm10 reference genome using STAR v2.2.1d. Gene counts were estimated using HTSeq (v0.6.1). Normalization and sample group comparisons of gene counts were performed using R package DESeq2 (v1.22.2). No filtering was performed prior to sample group comparisons, where the default DESeq2 independent filtering was applied. For the “Positive vs Negative” comparison a volcano plot was created, displaying significance and fold change for the dataset together with gene symbols for the most highly regulated genes. Heat map for validated genes found to differentially expressed (up- or down-regulated at least a 3-fold) between the two microglial populations of interest was generated using the Morpheus software (https://software.broadinstitute.org/Morpheus). The analysis of Gene Ontology (GO) terms was performed using DAVID^60^. Gene lists of interest were imported into DAVID and annotated regarding their biological processes (GOTERM_BP_FAT). Enrichment Map (www.baderlab.org/Software/EnrichmentMap)^61^, a plug in of cytoscape (www.cytoscape.org)^62^, was used as a visualization tool to produce a network graph of over-represented GO terms. Enrichment was based on FDR 0.05 and the jaccard coefficient of 0.5 was chosen to generate the network to determine the diversity and/or similarity of GO terms. Nodes represent enriched GO terms and node size correlates with the number of genes included in a specific GO term. Edges indicate the degree in overlap between nodes using the Jaccard coefficient. Only clusters ≥5 nodes were considered.

### Behaviour tests

The experimental mice were subjected to the subsequently described series of behavioural paradigms.

### Open Field

The open field test was used to asses both exploratory behaviour and locomotor activity. The mice were placed for 5 min in an open field (45×45×45cm3). Monitoring was done by an automated tracking system (SMART 2.5, Panlab). The behavioural parameters registered during 5-minute during sessions were the percentage of distance travelled in border and centre zones, and, to measure a possible anxiety behaviour, we calculated the time spent, in percentage, in border zone. *Arg1*^*Control*^ (n=10 females, n=8 males) and *Arg1*^*cKO*^ (n=9 females, n=6 males).

### Object Recognition Memory

The object recognition task was used for assessing recognition memory taking the advantage of the ability to discriminate the familiarity of previously met objects. Mice were tested as described previously^63^. Briefly, female mice were placed in a rectangular arena (45×45×45cm3) and two identical objects were placed in the arena during the training phase. Subsequently, one of the objects was substituted by novel one and the number of approximation to exploring the novel object as compared with the number of exploration of the familiar one assessed the animal’s memory. The relative exploration of the novel object was expressed as a discrimination index (#novel - #familiar)/(#novel + #familiar) taking account the training index. *Arg1*^*Control*^ (n=10) and *Arg1*^*cKO*^ (n=8).

### Y maze

Y maze is a test to investigate spatial memory. The maze was made of methacrylate and each arm was 18 cm long, 38 cm high, 8 cm wide and positioned at equal angles. Working memory was assessed by recording spontaneous exploring behaviour in a Y-maze. The mice were placed in the centre of maze allowing free exploration during 9 min. In this training, we measured the spontaneous alternation triplet and the total number of entries in each arms. To study spatial memory, we used the novel arm discrimination task based on the innate preference of rodents to explore a novel environment more than a familiar one. Therefore, we blocked a specific arm during 5 min. After 1 hour, the animals were placed again in the maze with all three arms opened allowed to explore the familiar arms and the novel one for 4 min. Percentage of entries in each arm in relation with the percentage of the first session were scored. The whole session was recorded by video and analysed later using SMART 2.5, Panlab. *Arg1*^*Control*^ (n=11 females, n=8 males) and *Arg1*^*cKO*^ (n=9 females, n=6 males).

### Rotarod

Rotarod paradigm was used to assess motor learning and neuromuscular coordination. To habituate mice to the rotarod (Ugo Basile Biological Research Apparatus), the animals were placed on the roller at a speed of 20 rpm until they could remain on it for one minute without falling off. To assay motor coordination mice were tested as described previously^64^. Briefly, animals were tested at a rotational speed of 20 rpm, accelerating to 60 rpm in increments of 5 rpm, during four successive trainings and quantifying the latency of the first fall and the number of total falls. Finally, the next day the protocol was repeated as a final test to estimate the motor memory in the all experimental mice groups. *Arg1*^*Control*^ (n=10 females, n=8 males) and *Arg1*^*cKO*^ (n=9 females, n=6 males).

### Statistical analysis

Statistical analysis was performed using GraphPad Prism (GraphPad Software, Version 6.0 or higher).

### Reporting summary

Further information is available in the Nature Research Reporting Summary linked to this article.

## Data availability

RNA-Seq data will be deposited in the Gene Expression Omnibus in the near future and definitely prior to article publication. All other data are available from the corresponding author upon reasonable request.

## Acknowledgements

We thank the Bioinformatics and Expression Analysis core facility, the Biomedicum Flow Cytometry core facility, the Biomedicum Imaging Core facility (with grants from the Strategic Research Area in Neuroscience (StratNeuro) and the Strategic Research Area in Stemc Cells and Regenerative Medicine (StratRegen) supported by the Swedish government) at the Karolinska Institutet for technical support. We would like to thank Sandra Vazquez for technical support. Supported by the Swedish Research Council and the Swedish Brain Foundation (B.J.), the Sigrid Jusélius Foundation and the Svenska Kulturfonden (V.S.), the Swedish Cancer Foundation (P.U. and B.J.), the Swedish Cancer Society (K.G., P.U. and B.J.), the Karolinska Institutet Foundation (P.G.R. and B.J.), the Wenner-Gren Foundation (K.G.), the Åke Wibergs Stiftelse (M.C.), the Spanish Ministerio de Ciencia, Innovación y Universidades/FEDER/UE RTI2018-098645-B-100 (J.L.V.), PID 2019-109569GB-100 (J.A.A.) and BFU2015-68655 (A.R.M.), the Spanish Junta de Andalucia /FEDER/EU P18-RT-1372 and the Spanish FEDER I+D+i-USE US-1264806 (J.L.V), the Swedish Childhood Cancer Foundation (K.B., L.K., P.U. and B.J.) and the Swedish governmental grants for researchers working in healthcare (K.B.), the Canada Research Chair (Tier 2) of Neuroimmune Plasticity in Health and Therapy to (M.E.T.), the TracInflam grant from ERA-NET NEURON Neuroinflammation (B.J., M.E.T. and M.H.).

## Contributions

V.S. performed the experiments except otherwise indicated; I.G.D., R.R. and J.L.V. created the *Arg1*^*cKO*^ mice, did the behavioural studies and analysed the generated data; S.K. and P.U. performed iDISCO and subsequent analysis; A.M.O. and K.B. performed tissue collection and processing and establishment of the cell isolation protocol; J.A.A. and E.M.P.V. did Golgi stainings and subsequent analysis; N.V. and M.E.T. performed and analysed the data of transmission electron microscopy; D.T. and M.T.H. did the morphometric analysis; K.G. and D.B. contributed in RNA-Seq data analysis; M.C. performed the RNA extraction and qPCR of the cell-sorted samples; M.C. and P.G.R. were involved in the quantification analysis; J.A.C. provided FACS sorting expertise; V.S. and B.J. initiated, conceptualised and designed the study, analysed and interpreted the data and wrote the first draft of the manuscript. All authors discussed the results and commented or edited the manuscript;

