## Supplementary Material for "*Arg1*^+^ microglia are critical for shaping cognition in female mice"

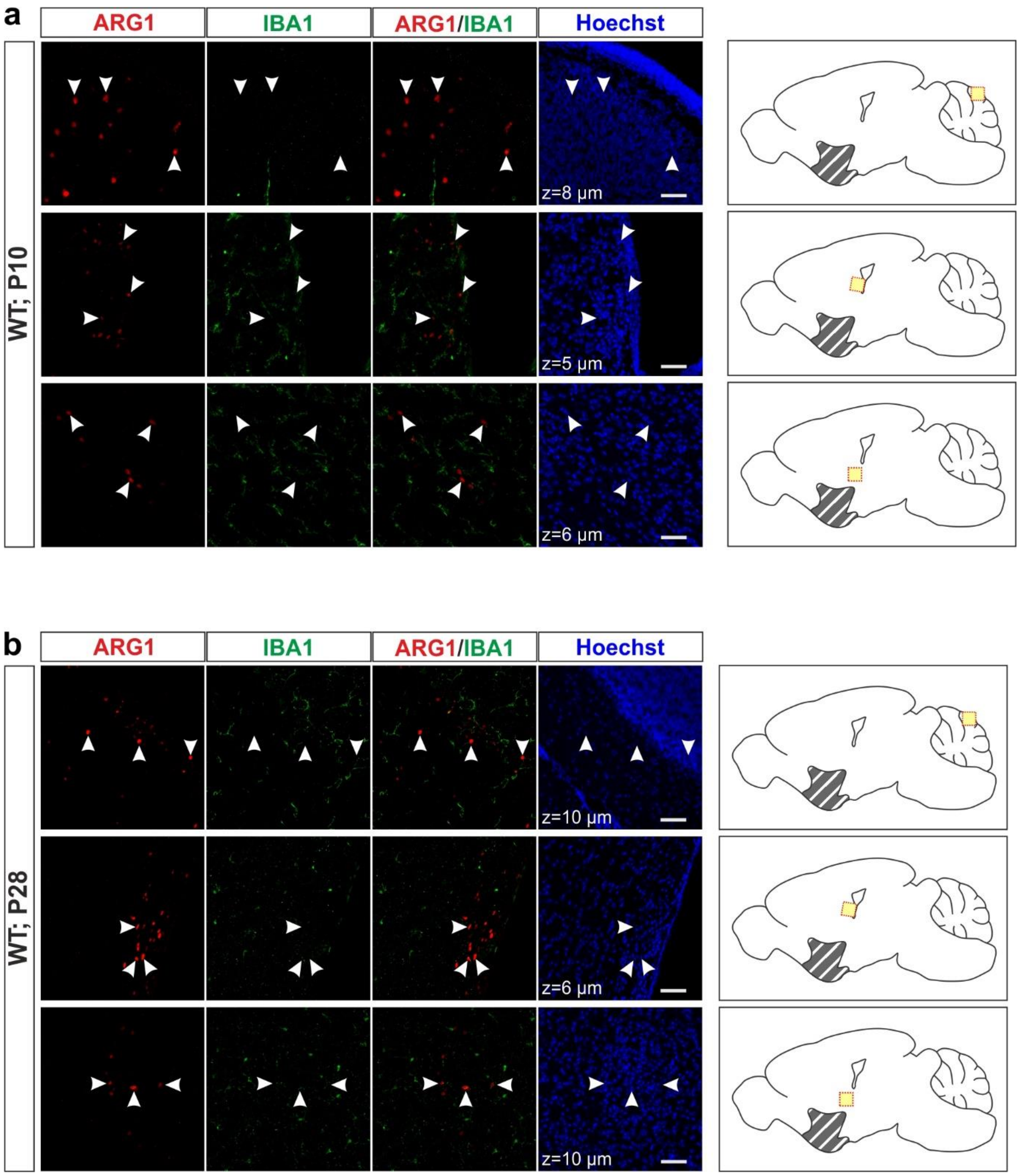

**Supplementary Figure 1**

**ARG1⁺ cells that are not microglia in WT mouse brains.**

**a-b**. $ARG1^+/IBA1^-$ cells in the cerebellum and around the ventricles (arrowheads) both at P10 (**a**) and at P28 (**b**). Scale bars, x=50 µm. Yellow squares indicate location of the corresponding images on their left; grey lines indicate the BF/vStr region.

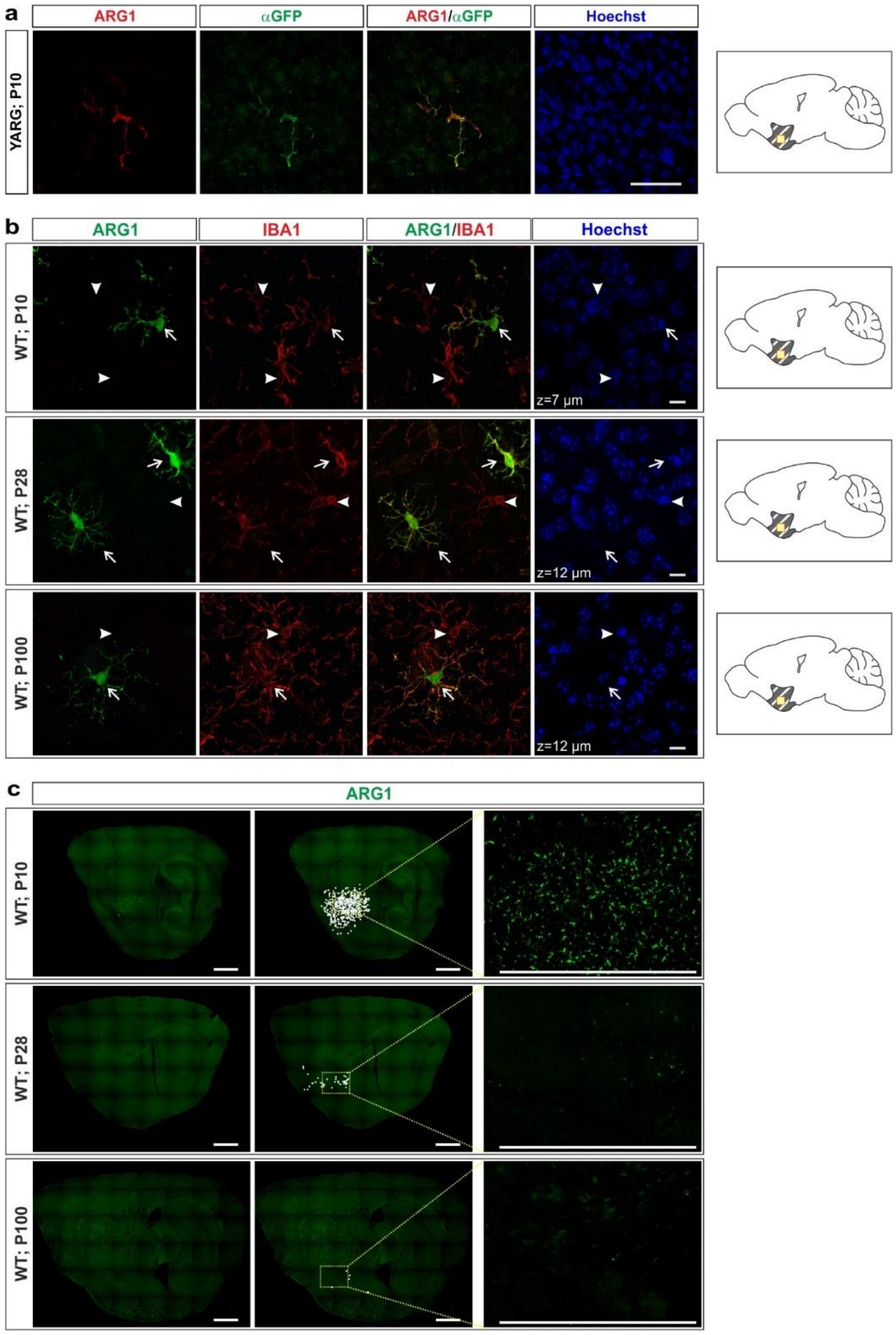

**Supplementary Figure 2**

***Arg1*⁺ microglia co-exist with "canonical" microglia in the same vicinity in WT mouse brain.**

**a**, *Arg1*-YFP⁺ microglia in male YARG mice, as recognised by α-GFP and α-ARG1 antibodies. **b**, *Arg1*⁺ microglia (arrows) and "canonical" *Arg1*⁻ microglia (arrowheads) during mouse brain development. **c**, *Arg1*⁺ microglia in BF/vStr at P10, P28 and P100. Each white dot represents a single *Arg1*⁺ microglia and has been manually annotated. Scale bars, x=50 µm, z=10 µm (**a**), x=10 µm (**b**), x=1000 µm, z=4 µm (**c**). Yellow squares indicate location of the corresponding images on their left; grey lines indicate the BF/vStr region.

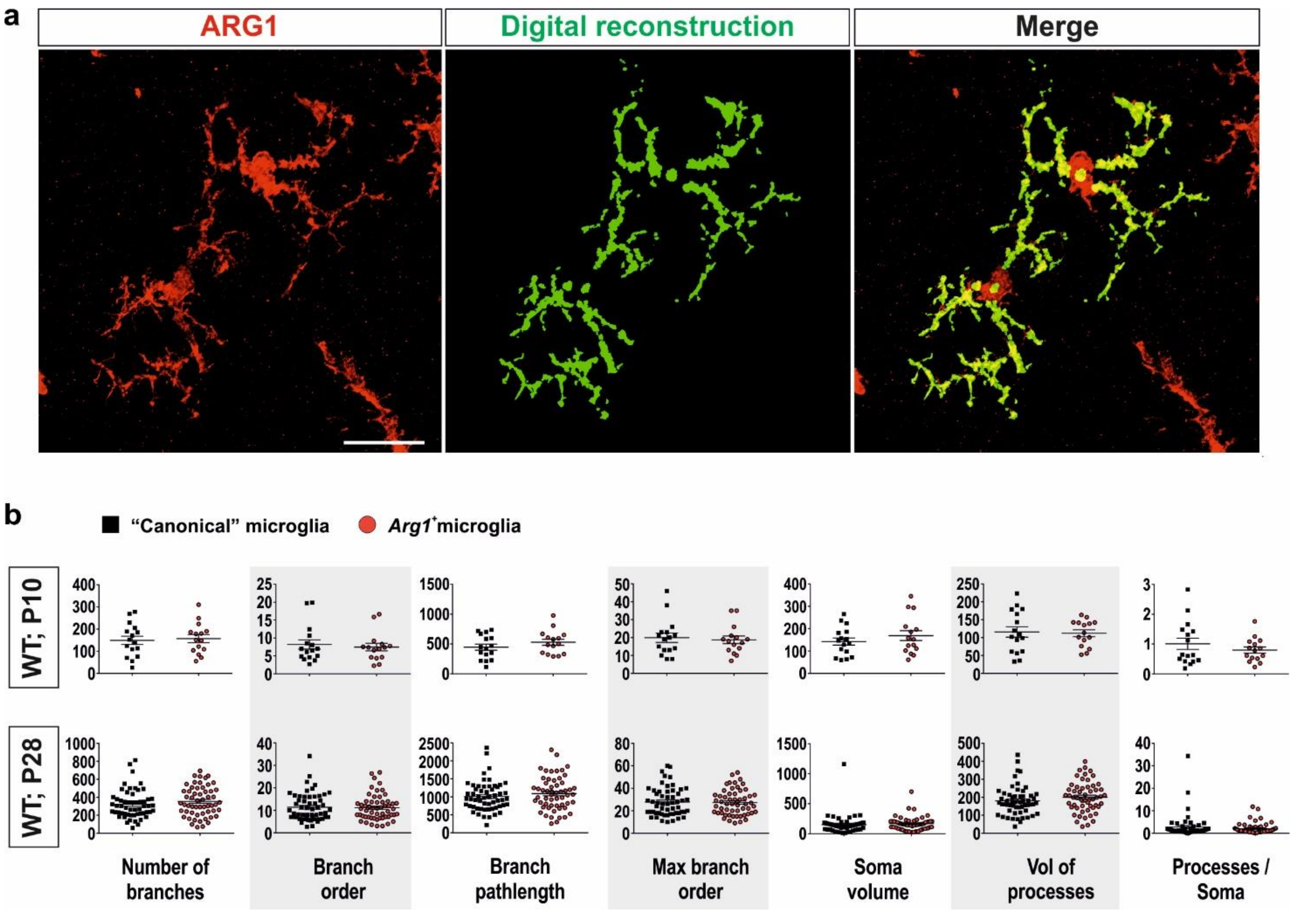

**Supplementary Figure 3**

**Morphometric comparison of "canonical" versus *Arg1*+microglia.**

**a**, Illustration of digital reconstruction of two *Arg1*+microglia. **b**, Comparison of *Arg1*+ microglia and "canonical" microglia from P10 (upper panel) and P28 (lower panel) mouse brain. Each square or circle represents a single microglial cell. Scale bar, x=10 µm, z=15 µm. Data in **b** are in mean ± s.e.m.

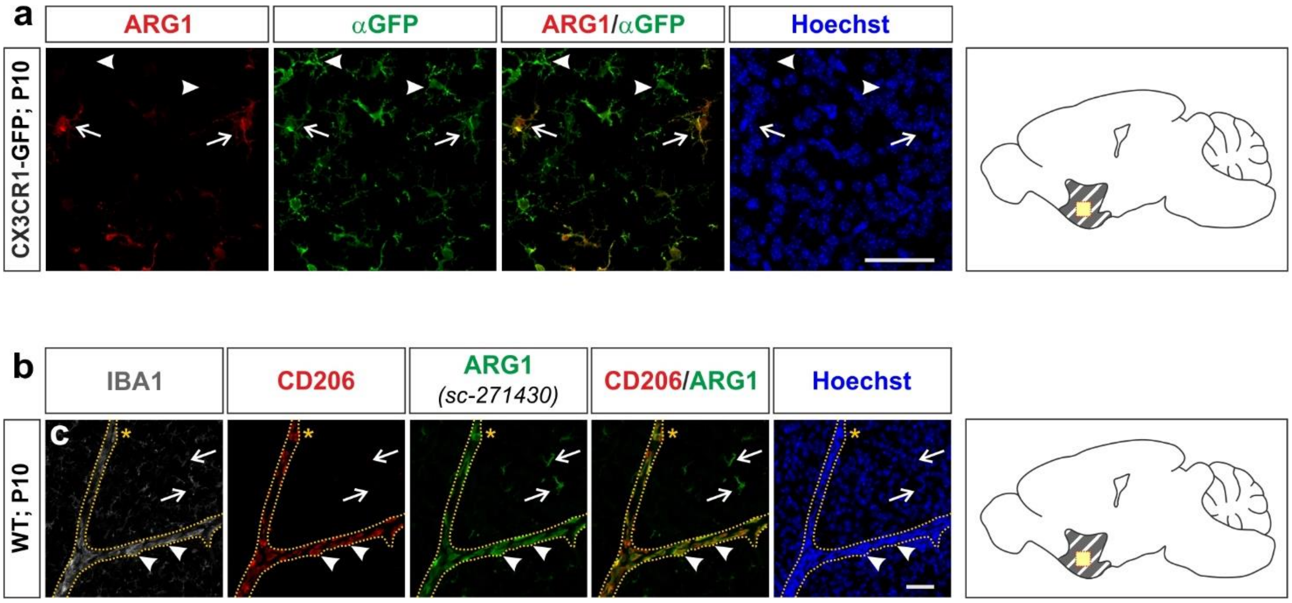

**Supplementary Figure 4**

***Arg1⁺* microglia co-localize with CX3CR1-GFP but not with CD206.**

**a**, CX3CR1-GFP$^+$/ARG1$^+$ microglia (arrows) coexist with CX3CR1-GFP$^+$/ARG1$^-$ microglia (arrowheads) in the BF/vStr of male animals. **b**, *Arg1*$^+$ microglia do not express the perivascular macrophage marker CD206 (arrows), while perivascular macrophages in this brain area express ARG1 (arrowheads). Note that *Arg1*$^+$ microglia are always ramified, while perivascular macrophages are amoeboid. Scale bars, x=50 µm, z=10 and 8.5 µm (for **a** and **b**, respectively). Yellow squares indicate location of the corresponding images on their left; grey lines indicate the BF/vStr region.

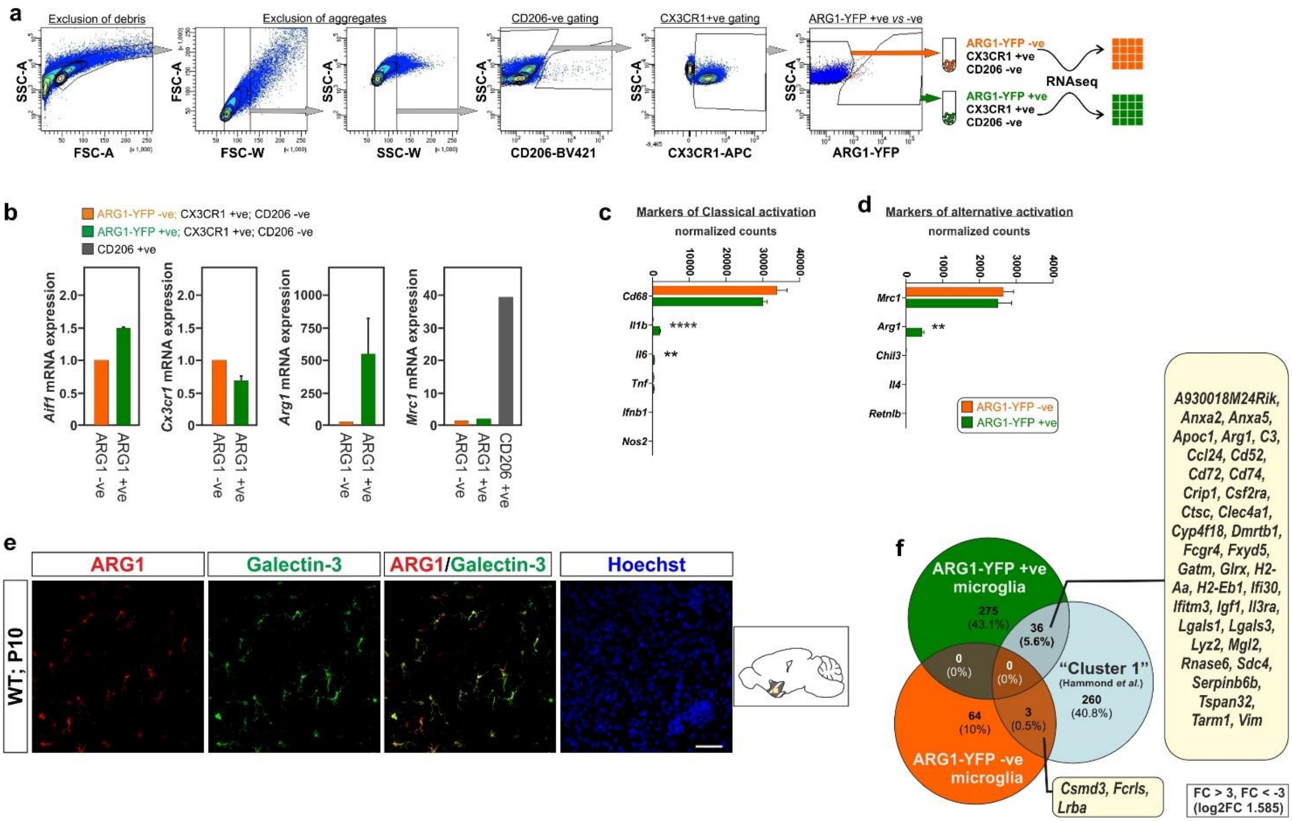

**Supplementary Figure 5**

***Arg1⁺*** **microglia express Galectin-3 and can not be classified as either "classical" or "alternative" activated microglia.**

**a**, Strategy for fluorescent-activated cell sorting of *Arg1-YFP*$^{+}$ and *Arg1-YFP*$^{-}$ microglia. **b**, Microglial mRNA expression determined by qPCR in "canonical" and *Arg1*$^{+}$ microglia (n=4-5 mice in 3 biological replicates (independent experiments) for *Aif1*, *Cx3cr1*, *Arg1*, and in 2 biological replicates for *Mrc1*) confirm successful isolation of the targeted cells. **c**-**d**, Although ARG1 is long been considered a marker of alternative activation, a number of alternative activation markers, namely *Mrc1, Chil3*, *Il4* and *Retnlb* are not expressed in *Arg1*$^{+}$ microglia higher than in "canonical" microglia (**d**). Same is true for classical activation markers, such as *Cd68, Tnf, Ifnb1* and *Nos2* (**c**). **e**, *Arg1*$^{+}$ microglia co-localize with the marker Galectin-3. Scale bar, x=50 µm, z=9 µm. **f**, Venn diagram showing overlaps between *Arg1*$^{+}$ microglia and "canonical" microglia presented here and "cluster 1" (reference [4]) (FC ≥3 or ≤ -3). Data in **b**-**d** are in mean ± s.e.m. Significant differences were determined by an unpaired two-tailed *t* test (**P*< 0.05, ***P*≤ 0.01, ****P*≤ 0.001, *****P*≤ 0.0001). Yellow squares indicate location of the corresponding images on their left; grey lines indicate the BF/vStr region.

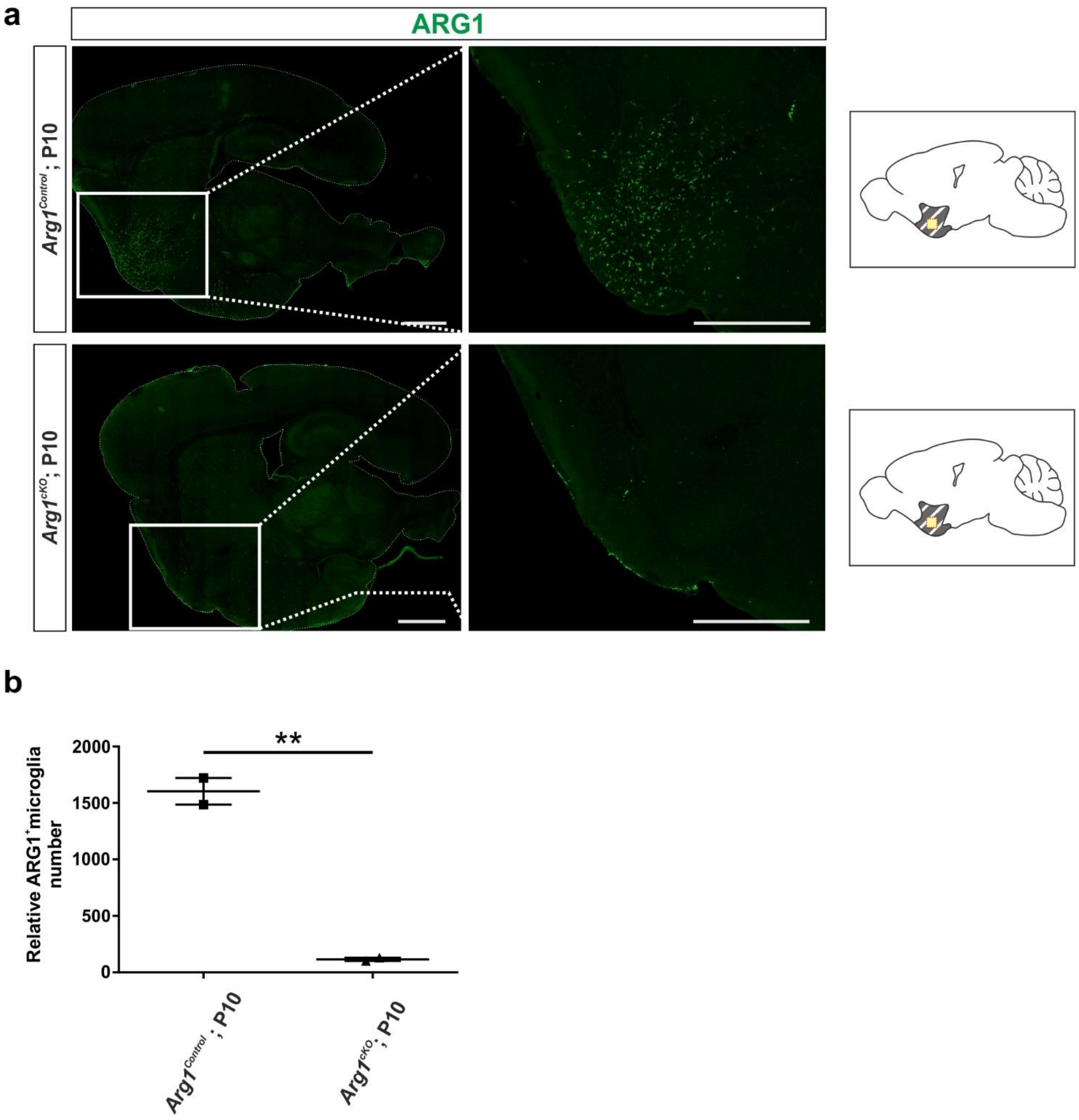

**Supplementary Figure 6**

***Arg1⁺* microglia are efficiently knocked down in *Arg1^cKO^*.**

**a**, Staining of matching sections shows that in *Arg1*$^{cKO}$ animals, only few *Arg1*$^{+}$ microglia remain when compared to control. **b**, Quantification of *Arg1*$^{+}$ microglia from matching sections (n=2 animals per genotype, 4 sections per animal). Scale bar, x=1000 µm. Data in **b** are in mean ± s.e.m. Significant differences were determined by an unpaired two-tailed *t* test (***P*≤ 0.01). Yellow squares indicate location of the corresponding images on their left; grey lines indicate the BF/vStr region.

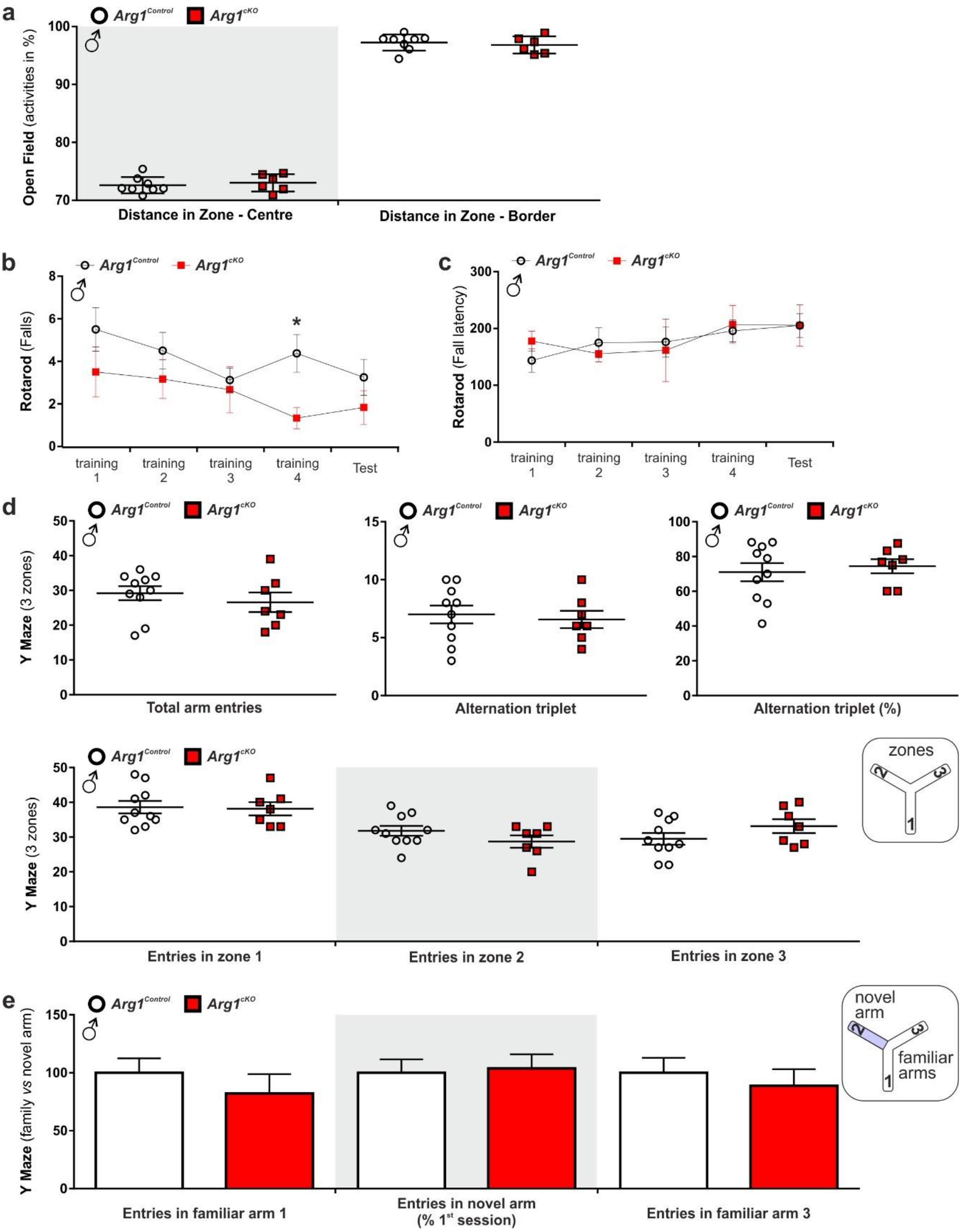

### Supplementary Figure 7

***Arg1$^{cKO}$* male animals do not show behavioural phenotype matching the female *Arg1$^{cKO}$* animals.**

**a**-**e**, *Arg1$^{cKO}$* male animals and controls were assessed for motoric (**a**-**c**) and memory (**d**-**e**) phenotypes. *Arg1$^{Control}$* n=8 (for **b**-**e**) and *Arg1$^{cKO}$* n=6 (for **b**-**e**). Data are in mean ± s.e.m. Significant differences were determined by an unpaired two-tailed *t* test (**P*< 0.05).

**Supplementary Table 1**

Quantification of *Arg1*$^+$ microglia clusters in P10 and P28 by iDISCO+ analysis (n=1 per developmental stage). Colour-code corresponds to clusters in Fig. 2a and Supplementary Movies 1-2.

| | Developmental stage | |
|---|---|---|
| | P10 | P28 |
| Loc1 | 17731 | 14217 |
| Loc2 | 315 | 280 |
| Loc3 | 968 | 319 |
| Loc4 | 23 | 44 |
| Loc5 | 1661 | 91 |
| Loc6 | 54 | NA |
| Loc7 | 88 | NA |
| Loc8 | 47 | NA |
| Loc9 | 150 | 16 |
| TOTAL | 21037 | 14967 |

### Supplementary Table 2

Registration of P28 *Arg1*$^+$ microglia from the BF/vStr to the Allen brain atlas (Developing mouse, P56), brain area acronyms (**a**), the area they occupy (**b**) and their absolute number (**c**).

(**a**)

| Acronym | Name |
|---|---|
| 7M | facial motor nucleus |
| AcbCo | accumbens nucleus, core domain |
| AcbSh | accumbens nucleus, shell domain |
| ACo | anterior cortical amygdaloid area |
| AHi | amygdalohippocampal area |
| ASt | amygdalo-striatal transition |
| BEC | bed nucleus of the external capsule |
| BLA | basolateral amygdaloid nucleus, anterior part |
| BMA | basomedial amygdaloid nucleus, anterior part |
| BMP | basomedial amygdaloid nucleus, posterior part |
| BSTLA | bed nucleus of stria terminalis, lateral amygdaloid division |
| BSTMC | bed nucleus of the stria terminalis, mediocentral division |
| CAT | nucleus of the central acoustic tract |
| CeC | central amygdalar nucleus, capsular part |
| CeL | central amygdalar nucleus, lateral part |
| CeM | central amygdaloid nucleus, medial part |
| Cl | claustrum |
| DEn | dorsal endopiriform nucleus |
| EPal | external globus pallidum |
| EPD | dorsal entopeduncular nucleus |
| HDB | horizontal nucleus of the diagonal band |
| HDBT | horizontal nucleus of the diagonal band, transitional part |
| IA | intercalated amygdaloid nuclei |
| ICjM | island of Calleja major |
| ICjPal | pallidal islands of Calleja |
| ICjStr | striatal islands of Calleja |
| InsC-6 | layer 6 of InsCx |
| IPAC | interstitial nucleus of the posterior limb of the anterior commissure |
| IPal | internal globus pallidus |
| isBLRt | isthmic part of basolateral isthmic reticular formation |
| isSNC | substantia nigra compacta, isthmic part |
| isSNR | substantia nigra reticulata, isthmic part |
| La | lateral amygdaloid nucleus |
| LOT | nucleus of the lateral olfactory tract |
| LOT1 | layer 1 of LOT |
| LOT2 | layer 2 of LOT |
| LOT3 | layer 3 of LOT |
| LPO1 | lateral preoptic area, PO1 part |
| LPO2 | lateral preoptic nucleus, PO2 part |
| LPrP1 | layer 1 of LPrP cortex |
| LPrP2 | layer 2 of LPrP cortex |
| LPrP3 | layer 3 of LPrP cortex |
| LSO | lateral superior olive |
| LSS | laterostriatal stripe |
| m1BRt | medial basal reticular formation of m1 |
| m1SNC | substantia nigra compacta, m1 part |
| m1SNR | substantia nigra reticulata, m1 part |
| m2BRt | reticular formation of basal m2 |
| m2SNC | m2 part of substantia nigra compacta |
| m2SNR | m2 part of substantia nigra reticulata |
| MCDB | magnocellular diagonal band nucleus |
| MeAD | medial amygdala, anterodorsal part |
| MeAV | medial amygdala, anteroventral part |
| MePD | medial amygdala, posterodorsal part |
| MePV | medial amygdala, posteroventral part |
| p1SNC | p1 part of the substantia nigra compacta |
| p1SNR | p1 part of the substantia nigra reticulata |
| p2SNC | p2 portion of the substantia nigra pars compacta |
| p2SNR | p2 portion of the substantia nigra pars reticulata |
| p3SNC | p3 portion of the substantia nigra pars compacta |
| p3SNR | p3 portion of the substantia nigra pars reticulata |
| PMCo | posteromedial cortical amygdaloid area |
| POHA | preopto-hypothalamic area |
| Put | putamen |
| r2-5Tr | r2 part of trigeminal transition zone |
| r2AVC | r2 part of anteroventral cochlear nucleus |
| r2SuVe | r2 part of superior vestibular nucleus |
| r3-5Tr | r3 part of trigeminal transition zone |
| r3AVC | r3 part of anteroventral cochlear nucleus |
| r3DPCRt | r3 part of dorsal parvicellular reticular formation |
| r3LVe | r3 part of lateral vestibular nucleus |
| r3Sp5O | r3 part of spinal trigeminal sensory column, oral part |
| r3VPCRt | r3 part of ventral parvicellular reticular formation |
| r4DPCRt | r4 part of dorsal parvicellular reticular formation |
| r4PVC | r4 part of posteroventral cochlear nucleus |
| r4Sp5O | r4 part of descending trigeminal sensory nucleus, oral part |
| r4VPCRt | r4 part of ventral parvicellular reticular formation |
| r5DPCRt | r5 part of dorsal parvicellular reticular formation |
| r5Sp5O | r5 part of the oral Sp5 subnucleus |
| r5SpVe | r5 part of spinal vestibular nucleus |
| r6-5Tr | r6 part of trigeminal transition zone |
| r6DPCRt | r6 part of dorsal parvicellular reticular formation |
| r6Sp5O | r6 part of spinal trigeminal nucleus, pars oralis |
| r6VPCRt | r6 part of ventral parvicellular reticular formation |
| r7-5Tr | r7 part of trigeminal transition zone |
| r7Sp5I | r7 part of spinal trigeminal nucleus, interpolar part |
| r7VPCRt | r7 part of ventral parvicellular reticular formation |
| r8Sp5I | r8 part of spinal trigeminal nucleus, interpolar part |
| r9Sp5I | r9 part of spinal trigeminal nucleus, interpolar part |
| Rt | reticular nucleus |
| SePalCo | septopallidal core nucleus |
| SePalSh | septopallidal shell area |
| SIB | substantia innominata/basal nucleus |
| SIBT | substantia innominata/basal nucleus, transitional part |
| SM | nucleus of the stria medullaris (prethalamic) |
| TuPal1 | plexiform layer of TuPal |
| TuPal2 | corticoid layer of TuPal |
| TuPal3 | polymorph layer of TuPal |
| TuSePal | septopallidal part of the olfactory tuberculum |
| TuSeStr | septostriatal part of the olfactory tuberculum |
| TuStr1 | plexiform layer of TuStr |
| TuStr2 | corticoid layer of TuStr |
| TuStr3 | polymorph layer of TuStr |
| UDLH | upper dorsal lateral hypothalamic area |
| UDPeF | upper dorsal perifornical nucleus |
| VEn | ventral endopiriform nucleus |
| VLPeO | ventrolateral periolivary nucleus |
| VPal | ventral pallidum |
| VPrP1 | layer 1 of VPrP cortex |
| VPrP2 | layer 2 of VPrP cortex |
| VPrP3 | layer 3 of VPrP cortex |
| VStr | ventral striatum |

(**b**)

| Acronym | Percentage #1 | Percentage #2 | Percentage #3 | Average |
|---|---|---|---|---|
| VPal | 100 | 99.07255164 | 99.98843901 | 99.68699688 |
| CeM | 99.69814575 | 100 | 99.18068133 | 99.62627569 |
| MCDB | 99.92016322 | 100 | 98.52745498 | 99.4825394 |
| LSS | 100 | 98.0717729 | 97.42903053 | 98.50026781 |
| TuPal3 | 99.61554192 | 93.89366053 | 98.52351738 | 97.34423995 |
| ICjPal | 96.46347138 | 94.78827362 | 94.78827362 | 95.34667287 |
| TuPal2 | 95.75583157 | 89.50015147 | 93.62011512 | 92.95869938 |
| VStr | 99.23411644 | 87.40374776 | 86.80427076 | 91.14737832 |
| IPAC | 83.85151018 | 90.7792611 | 90.30913204 | 88.3133011 |
| LOT | 81.1452514 | 96.62476723 | 80.86592179 | 86.21198014 |
| SIB | 90.80228301 | 86.02412233 | 77.94744777 | 84.9246177 |
| TuStr3 | 86.16847246 | 69.38760995 | 66.60186349 | 74.05264863 |
| HDB | 79.23211475 | 74.75774007 | 64.10355081 | 72.69780188 |
| TuStr2 | 80.78417675 | 62.17089464 | 61.40243636 | 68.11916925 |
| TuPal1 | 64.37714356 | 57.48713866 | 55.33133268 | 59.06520496 |
| CeC | 50.38705137 | 60.88142153 | 51.84729064 | 54.37192118 |
| DEn | 43.20618684 | 53.14887848 | 45.67722455 | 47.34409662 |
| ICjStr | 55.29122231 | 42.49384742 | 43.56029532 | 47.11512168 |
| LOT3 | 58.8080631 | 53.88548057 | 28.10400234 | 46.93251534 |
| VEn | 47.65077091 | 48.72505169 | 41.32940556 | 45.90174272 |
| ASt | 40.49874417 | 51.34553283 | 40.64585576 | 44.16337759 |
| CeL | 43.79373076 | 49.20235097 | 37.22362161 | 43.40656778 |
| VPrP3 | 43.35215284 | 44.42429427 | 36.19047619 | 41.32230777 |
| IPal | 44.03290859 | 35.38556714 | 29.03202481 | 36.15016685 |
| TuStr1 | 48.33526012 | 30.26011561 | 29.45086705 | 36.01541426 |
| VPrP2 | 35.48321835 | 39.50818346 | 27.81363167 | 34.26834449 |
| Put | 37.20535278 | 32.64229342 | 30.94016147 | 33.59593589 |
| EPal | 37.85915314 | 28.66016865 | 25.41328118 | 30.64420099 |
| CSPall | 27.5882552 | 27.86608519 | 27.62547464 | 27.69327168 |
| BEC | 32.77145124 | 22.8763216 | 22.25224646 | 25.9666731 |
| SePalSh | 43.16017724 | 4.907136796 | 14.76854907 | 20.9452877 |
| SePalCo | 43.00958628 | 5.713925328 | 13.67305752 | 20.79885637 |
| SIBT | 41.21827605 | 5.935142139 | 14.33008155 | 20.49449991 |
| MeAD | 22.93920179 | 23.48004476 | 12.58112645 | 19.666791 |
| TuSePal | 30.54626987 | 3.456991439 | 22.80065226 | 18.93463786 |

| Acronym | Percentage #1 | Percentage #2 | Percentage #3 | Average |
|---|---|---|---|---|
| VPrP1 | 19.68087855 | 20.46511628 | 14.10271318 | 18.08290267 |
| UDLH | 14.94654412 | 24.1114133 | 13.01697459 | 17.35831067 |
| VPall | 15.78912078 | 16.60011734 | 13.58645545 | 15.32523119 |
| LOT2 | 15.97851359 | 18.54405516 | 4.01667602 | 12.84641492 |
| ASPall | 10.72913754 | 14.58094662 | 10.63171473 | 11.98059963 |
| BMA | 9.93658029 | 11.24213447 | 9.002625972 | 10.06044691 |
| LPO1 | 18.70059577 | 7.236867723 | 4.07110315 | 10.00285555 |
| PaSe | 16.33096024 | 2.092377149 | 3.959020663 | 7.460786016 |
| BLA | 1.802870359 | 9.435416914 | 10.73716048 | 7.325149251 |
| Cl | 11.36119488 | 4.199293323 | 3.441492047 | 6.333993416 |
| BSTLA | 1.552170164 | 11.59816039 | 4.943949411 | 6.031426655 |
| HDBT | 10.70822491 | 1.173775314 | 5.817965833 | 5.899988687 |
| AcbSh | 16.10491301 | 0 | 0.320861912 | 5.475258309 |
| AcbCo | 14.52543984 | 0.584158825 | 0.913437031 | 5.341011899 |
| TuSeStr | 12.20408647 | 0 | 0.015808402 | 4.073298291 |
| La | 2.382703564 | 4.800649689 | 3.496674941 | 3.560009398 |
| IA | 0.16634981 | 3.814163498 | 6.202471483 | 3.394328264 |
| LPrP3 | 4.006919649 | 3.403580833 | 1.313199315 | 2.907899932 |
| LPO2 | 3.917525773 | 3.223992502 | 0.987191503 | 2.709569926 |
| LPall | 2.962487825 | 3.322235458 | 1.548951132 | 2.611224805 |
| LPrP2 | 2.285117629 | 3.585222968 | 1.325978082 | 2.398772893 |
| ICjM | 6.765861875 | 0 | 0 | 2.255287292 |
| EPD | 1.369948705 | 3.878393594 | 0.537970724 | 1.928771008 |
| LOT1 | 1.857585139 | 2.850432369 | 0.80068325 | 1.836233586 |
| POA | 2.056675822 | 1.782005761 | 0.864205802 | 1.567629128 |
| p3A | 0.491858586 | 2.773585241 | 1.104462241 | 1.456635356 |
| LPrP1 | 0.798948101 | 1.473484357 | 0.583143669 | 0.951858709 |
| PHyA | 0.830395097 | 1.052822355 | 0.583253699 | 0.82215705 |
| TelA | 1.006347031 | 0.830277662 | 0.62756278 | 0.821395824 |
| POHA | 0.629249975 | 0.781487872 | 0.005074597 | 0.471937481 |
| MeAV | 0.608642727 | 0.311929397 | 0 | 0.306857375 |
| ACo | 0.081092883 | 0.224564905 | 0.024951656 | 0.110203148 |
| THyA | 0.013277442 | 0.013277442 | 0 | 0.008851628 |
| InsC-6 | 0.025423056 | 0 | 0 | 0.008474352 |

(c)

| Acronym | | Brain | | | | Average |
|---|---|---|---|---|---|---|
| | | #1 | #2 | #3 | | |
| Put | | 2870 | 3349 | 2974 | | 3064.333333 |
| VStr | | 1606 | 1580 | 1696 | | 1627.333333 |
| VPal | | 1410 | 1365 | 1321 | | 1365.333333 |
| TuPal3 | | 1037 | 1095 | 1104 | | 1078.666667 |
| TuPal2 | | 836 | 872 | 1029 | | 912.3333333 |
| TuPal1 | | 604 | 700 | 682 | | 662 |
| VPrP2 | | 564 | 639 | 374 | | 525.6666667 |
| TuStr3 | | 502 | 433 | 550 | | 495 |
| SIB | | 491 | 495 | 382 | | 456 |
| EPal | | 503 | 379 | 376 | | 419.3333333 |
| VPrP1 | | 397 | 498 | 350 | | 415 |
| HDB | | 354 | 411 | 360 | | 375 |
| VPrP3 | | 349 | 415 | 311 | | 358.3333333 |
| TuStr2 | | 374 | 273 | 408 | | 351.6666667 |
| IPal | | 372 | 291 | 328 | | 330.3333333 |
| VEn | | 173 | 374 | 176 | | 241 |
| TuStr1 | | 253 | 196 | 262 | | 237 |
| IPAC | | 191 | 293 | 198 | | 227.3333333 |
| MCDB | | 141 | 206 | 112 | | 153 |
| DEn | | 86 | 179 | 75 | | 113.3333333 |
| BEC | | 78 | 91 | 94 | | 87.66666667 |
| p2SNR | | 82 | 69 | 109 | | 86.66666667 |
| r5Sp5O | | 61 | 82 | 91 | | 78 |
| p1SNR | | 74 | 79 | 77 | | 76.66666667 |
| MePV | | 55 | 33 | 67 | | 51.66666667 |
| r6Sp5O | | 33 | 53 | 60 | | 48.66666667 |
| Cl | | 53 | 47 | 36 | | 45.33333333 |
| r5DPCRt | | 44 | 29 | 54 | | 42.33333333 |
| CeL | | 38 | 47 | 37 | | 40.66666667 |
| TuSePal | | 38 | 5 | 74 | | 39 |
| ICjPal | | 38 | 40 | 37 | | 38.33333333 |
| LOT | | 36 | 35 | 32 | | 34.33333333 |
| CeM | | 42 | 31 | 27 | | 33.33333333 |
| UDLH | | 22 | 51 | 27 | | 33.33333333 |
| r4Sp5O | | 23 | 24 | 46 | | 31 |
| CeC | | 24 | 33 | 33 | | 30 |
| p1SNC | | 24 | 23 | 41 | | 29.33333333 |
| LSS | | 41 | 25 | 20 | | 28.66666667 |
| ASt | | 25 | 35 | 20 | | 26.66666667 |
| LPrP2 | | 12 | 43 | 23 | | 26 |
| r7Sp5I | | 22 | 22 | 29 | | 24.33333333 |
| p3SNR | | 16 | 10 | 40 | | 22 |
| SIBT | | 37 | 20 | 9 | | 22 |
| r8Sp5I | | 16 | 18 | 26 | | 20 |
| LPrP1 | | 14 | 31 | 14 | | 19.66666667 |
| ICjStr | | 10 | 23 | 22 | | 18.33333333 |
| r4DPCRt | | 21 | 17 | 13 | | 17 |
| r3Sp5O | | 11 | 23 | 13 | | 15.66666667 |
| m1SNC | | 3 | 13 | 21 | | 12.33333333 |
| MePD | | 4 | 12 | 17 | | 11 |
| SePalSh | | 23 | 0 | 8 | | 10.33333333 |
| SePalCo | | 20 | 7 | 1 | | 9.333333333 |
| m1BRt | | 5 | 8 | 14 | | 9 |
| r6DPCRt | | 16 | 4 | 6 | | 8.666666667 |
| HDBT | | 4 | 8 | 9 | | 7 |
| r6-5Tr | | 5 | 9 | 6 | | 6.666666667 |
| BSTMC | | 4 | 7 | 7 | | 6 |

| Acronym | | Brain | | | | Average |
|---|---|---|---|---|---|---|
| | | #1 | #2 | #3 | | |
| p2SNC | | 2 | 5 | 11 | | 6 |
| r4VPCRt | | 12 | 3 | 3 | | 6 |
| isSNC | | 2 | 10 | 5 | | 5.666666667 |
| m1SNR | | 7 | 6 | 4 | | 5.666666667 |
| r3AVC | | 8 | 6 | 3 | | 5.666666667 |
| LPrP3 | | 4 | 7 | 5 | | 5.333333333 |
| Rt | | 9 | 4 | 3 | | 5.333333333 |
| BLA | | 1 | 10 | 4 | | 5 |
| LSO | | 7 | 5 | 2 | | 4.666666667 |
| MeAV | | 5 | 2 | 7 | | 4.666666667 |
| isBLRt | | 2 | 7 | 4 | | 4.333333333 |
| La | | 2 | 7 | 4 | | 4.333333333 |
| LPO1 | | 11 | 1 | 0 | | 4 |
| AcbCo | | 8 | 1 | 2 | | 3.666666667 |
| r7-5Tr | | 6 | 3 | 2 | | 3.666666667 |
| CAT | | 6 | 2 | 1 | | 3 |
| m2SNR | | 2 | 2 | 5 | | 3 |
| MeAD | | 1 | 0 | 6 | | 2.333333333 |
| r3VPCRt | | 5 | 1 | 0 | | 2 |
| UDPeF | | 1 | 4 | 1 | | 2 |
| LOT3 | | 0 | 6 | 0 | | 2 |
| m2SNC | | 0 | 0 | 6 | | 2 |
| r2A | | 0 | 6 | 0 | | 2 |
| AHi | | 0 | 3 | 2 | | 1.666666667 |
| p3A | | 0 | 3 | 2 | | 1.666666667 |
| AcbSh | | 4 | 0 | 0 | | 1.333333333 |
| r2AVC | | 3 | 1 | 0 | | 1.333333333 |
| isSNR | | 0 | 2 | 2 | | 1.333333333 |
| LOT2 | | 0 | 3 | 1 | | 1.333333333 |
| m1B | | 0 | 2 | 2 | | 1.333333333 |
| p3SNC | | 0 | 0 | 4 | | 1.333333333 |
| LPO2 | | 3 | 0 | 0 | | 1 |
| r3DPCRt | | 2 | 1 | 0 | | 1 |
| BMP | | 0 | 0 | 3 | | 1 |
| r3-5Tr | | 0 | 3 | 0 | | 1 |
| r3LVe | | 0 | 2 | 1 | | 1 |
| r5SpVe | | 0 | 1 | 2 | | 1 |
| r6VPCRt | | 0 | 0 | 3 | | 1 |
| m2BRt | | 1 | 1 | 0 | | 0.666666667 |
| r7VPCRt | | 1 | 0 | 1 | | 0.666666667 |
| VLPeO | | 2 | 0 | 0 | | 0.666666667 |
| BMA | | 0 | 0 | 2 | | 0.666666667 |
| BSTLA | | 0 | 1 | 1 | | 0.666666667 |
| isB | | 0 | 1 | 1 | | 0.666666667 |
| r2-5Tr | | 0 | 2 | 0 | | 0.666666667 |
| InsC-6 | | 1 | 0 | 0 | | 0.333333333 |
| r4PVC | | 1 | 0 | 0 | | 0.333333333 |
| r9Sp5I | | 1 | 0 | 0 | | 0.333333333 |
| TuSeStr | | 1 | 0 | 0 | | 0.333333333 |
| 7M | | 0 | 0 | 1 | | 0.333333333 |
| EPD | | 0 | 1 | 0 | | 0.333333333 |
| LOT1 | | 0 | 1 | 0 | | 0.333333333 |
| PMCo | | 0 | 1 | 0 | | 0.333333333 |
| r3-5Tr | | 0 | 0 | 1 | | 0.333333333 |
| r2SuVe | | 0 | 1 | 0 | | 0.333333333 |
| SM | | 0 | 1 | 0 | | 0.333333333 |
